# Activity-Based Ratiometric FRET Probe Reveals Oncogene-Driven Changes in Labile Copper Pools Induced by Altered Glutathione Metabolism

**DOI:** 10.1101/682047

**Authors:** Clive Yik-Sham Chung, Jessica M. Posimo, Sumin Lee, Tiffany Tsang, Julianne M. Davis, Donita C. Brady, Christopher J. Chang

## Abstract

Copper is essential for life, and beyond its well-established ability to serve as a tightly-bound, redox-active active site cofactor for enzyme function, emerging data suggest that cellular copper also exists in labile pools, defined as loosely bound to low molecular weight ligands, which can regulate diverse transition metal signaling processes spanning neural communication and olfaction, lipolysis, rest-activity cycles, and kinase pathways critical for oncogenic signaling. To help decipher this growing biology, we report a first-generation ratiometric fluorescence resonance energy transfer (FRET) copper probe, FCP-1, for activity-based sensing of labile Cu(I) pools in live cells. FCP-1 links fluorescein and rhodamine dyes through a tris[(2-pyridyl)methyl]amine (TPA) bridge. Bioinspired Cu(I)-induced oxidative cleavage decreases FRET between fluorescein donor and rhodamine acceptor. FCP-1 responds to Cu(I) with high metal selectivity and oxidation-state specificity and facilitates ratiometric measurements that minimize potential interferences arising from variations in sample thickness, dye concentration, and light intensity. FCP-1 enables imaging of dynamic changes in labile Cu(I) pools in live cells in response to copper supplementation/depletion, differential expression of the copper importer CTR1, and redox stress induced by manipulating intracellular glutathione levels and GSH/GSSG ratios. FCP-1 imaging reveals a labile Cu(I) deficiency induced by oncogene-driven cellular transformation that promotes fluctuations in glutathione metabolism, where lower GSH/GSSG ratios decrease labile Cu(I) availability without affecting total copper levels. By connecting copper dysregulation and glutathione stress in cancer, this work provides a valuable starting point to study broader crosstalk between metal and redox pathways in health and disease with activity-based probes.

**Significance:** Copper is a required metal nutrient for life, yet its altered homeostasis is associated with many diseases. Thus, to develop new methods to help decipher copper biology, we present an activity-based ratiometric FRET probe that exploits a biomimetic, copper(I)-dependent cleavage reaction to enable imaging of loosely-bound, labile copper pools in cells with metal and oxidation state selectivity and a self-calibrating ratiometric response. Application of this technology to cellular models of cancer reveals that oncogene-driven changes in the metabolism of glutathione, a major cellular redox buffer, leads to a labile copper(I) deficiency. This work establishes the relevance of copper dysregulation to cancer metabolism and presages further opportunities for activity-based sensing in studies of metal biology.

## Introduction

Copper is a required redox-active nutrient for life (1), serving as a potent catalytic and/or structural cofactor in proteins for fundamental processes such as oxygen transport, respiration and metabolism, cell growth and differentiation, and signal transduction (2–6). In addition to tightly-bound copper pools buried within metalloprotein active sites, emerging data suggest the presence of labile copper pools that are bound relatively weakly by low molecular weight ligands (7). Labile copper pools contribute to a growing number of dynamic transition metal signaling pathways (8) including neural communication (9–11), olfaction (12, 13), lipolysis (6), rest-activity cycles (14), and kinase pathways involved in signal transduction and oncogenesis (5, 15). Indeed, copper dysregulation can result in aberrant oxidative and nitrosative stress events that accompany diseases spanning cancer (5, 16), cardiovascular disorders, neurodegenerative Alzheimer’s, Parkinson’s and Huntington’s diseases (3, 17), diabetes and obesity, and genetic Menkes and Wilson disorders (18–21).

The signal/stress dichotomy of copper motivates the development of new chemical tools to help decipher its broader contributions to physiology and pathology. In this context, fluorescent (6, 10, 11, 14, 22–38), bioluminescent (39), and magnetic resonance imaging probes (40–42) for visualizing labile copper pools with metal and oxidation state specificity, particularly when used in concert with complementary techniques for measuring total metal content (22, 43, 44), including laser ablation inductively-coupled plasma mass spectrometry (LA-ICP-MS) (45–47), nanoscale secondary ion mass spectrometry (nano-SIMS) (38, 48), and X-ray fluorescence microscopy (XFM) (10, 49, 50), can provide a coherent picture of metal homeostasis for a given biological model. Along these lines, a chemical limitation in fluorescent probe development is that the vast majority of copper indicators to date only respond through changes in fluorescence intensity in a single color, which can complicate quantitative measurements of copper pools and/or comparison of copper levels across different biological specimens owing to potential variations in probe uptake along with interferences arising from copper-independent phenomena, such as heterogeneities in sample thickness and light intensity (51, 52). To address this issue, ratiometric probes, which utilize excitation/emission spectral changes over two or more wavelengths, can in principle enable internal self-calibration for non-homogenous probe loading and limit interferences across variable specimens (53–57). However, the few ratiometric fluorescent copper indicators reported to date are limited largely by a lack of oxidation state specificity and/or decrease in overall fluorescence intensity upon copper detection (31–37).

We now present the design, synthesis, and biological applications of a first-generation ratiometric fluorescent copper indicator, FRET Copper Probe 1 (FCP-1). This activity-based sensing probe exploits the chemoselective Cu(I)-dependent oxidative C–O cleavage of a tris[(2-pyridyl)methyl]amine (TPA) (28, 58, 59) unit linking fluorescein donor and rhodamine acceptor units (Scheme 1). The high metal and oxidation state specificity of FCP-1 for Cu(I), along with its two-color ratiometric FRET response, enables its use to monitor dynamic changes in labile Cu(I) pools in live cells. Owing to the self-calibration afforded by its ratiometric response, FCP-1 was capable of reliably detecting decreases in endogenous levels of labile Cu(I) in mouse embryonic fibroblasts (MEFs) lacking the high-affinity copper transporter *Ctr1* (*Ctr1*^*−/−*^) relative to wildtype congeners. Further experiments identified fluctuations in labile Cu(I) levels, but not total copper levels, under redox stress induced by genetic alterations to glutathione biosynthesis pathways. Interestingly, a deficiency in labile Cu(I) pools was found by introduction of oncogenic BRAF^V600E^ or KRAS^G12D^, linking labile copper dysregulation to oncogene-dependent redox status. The ability of FCP-1 to monitor labile Cu(I) pools with a ratiometric response provides a starting point for further studies of redox contributions to transition metal signaling.

## Results and Discussion

### Design, Synthesis and Characterization of FCP-1

Inspired by bioinorganic TPA model compounds that exhibit Cu(I)-dependent oxidative cleavage chemistry (28, 58, 59), our design of FCP-1 in Scheme 1 connects a fluorescein FRET donor and rhodamine FRET acceptor with copper-responsive TPA linker. Changes in labile Cu(I) levels would modulate intramolecular FRET efficiency and lead to different fluorescence intensities of the donor and acceptor at two different wavelengths that can be self-calibrated in a ratiometric manner. Under lower Cu(I) levels, the probe stays intact with higher FRET and rhodamine-dominant emission, with higher Cu(I) levels leading to copper-dependent oxidative cleavage of the TPA linker and lower FRET and fluorescein-dominant emission.

The synthesis of FCP-1 starts with preparation of the 2-bromomethylpyridine-functionalized fluorescein methyl ester (**2**) and picolylamine-functionalized rhodamine (**5**; Scheme 2). Compound **2** was synthesized in 2 steps from the previously reported fluorescein methyl ester by nucleophilic substitution onto 6-(bromomethyl)-2-pyridinemethanol and subsequent bromination of the methyl alcohol. In parallel, nosyl-protected picolylamine (**3**) was prepared by monosubstitution of bis(chloromethyl)pyridine from 4-nitro-*N*-(2-pyridinylmethyl)benzenesulfonamide and then reacted with piperazinyl-functionalized rhodamine (Pz-rhodamine) to afford **4**. Subsequently nosyl group of **4** is deprotected by thiophenol and K_2_CO_3_ to yield **5**. With the key pieces **2** and **5** in hand, TPA motif can be generated through nucleophilic substitution of 2-bromomethyl group on **2** by the picolylamine on **5** under basic conditions, furnishing the final ratiometric probe FCP-1 that was characterized by ^1^H and ^13^C{^1^H} NMR, high-resolution mass spectrometry (HRMS), and liquid chromatography-coupled mass spectrometry (LC−MS; Fig. S1). The intensity-based Cu(I) probe FL-TPA was prepared as a control compound with the same Cu(I)-dependent activity-based sensing trigger and fluorescein scaffold but lacking a ratiometric response (Scheme 2).

### Reactivity and Selectivity of FCP-1 to Copper

The intracellular environment is known to be reducing and contains high concentrations of biomolecules that results in macromolecular crowding (60, 61). As such, to mimic aspects of these properties *in vitro*, we evaluated FCP-1 in PBS buffer containing glutathione (2 mM) and PEG-400 (40%, v/v). FCP-1 showed absorption maxima at 465, 493 and 552 nm, with molar extinction coefficients of 20,000, 22,100 and 40,000 M^−1^ cm^−1^, respectively (Fig. S2). The first two absorption maxima corresponded to the absorption of the fluorescein moiety, as similar absorption maxima at 460 and 489 nm were found in the absorption spectra of FL-TPA (Fig. S2) and reported monocaged fluorescein methyl ester (62, 63), whereas the lowest energy absorption at 552 nm originated from the rhodamine moiety (Fig. S2). Upon photoexcitation at 458 nm, FCP-1 was found to emit at 526 and 576 nm (Fig. 1A), corresponding to fluorescein and rhodamine signatures, respectively. Notably, the Pz-rhodamine building block shows only weak absorption at 458 nm (Fig. S2). Together with the significant contribution of the lower-energy fluorescence from the excitation at 450–500 nm, which is in the region of FL-TPA absorption (Fig. S3), the data support the occurence of FRET from the fluorescein donor to the rhodamine acceptor. Using FL-TPA as a reference FRET donor compound (Fig. S4) with assumption that quenching of donor fluorescence originated only from the FRET process, the FRET efficiency in the FCP-1 cassette is estimated to be 84%.

**Fig. 1.**
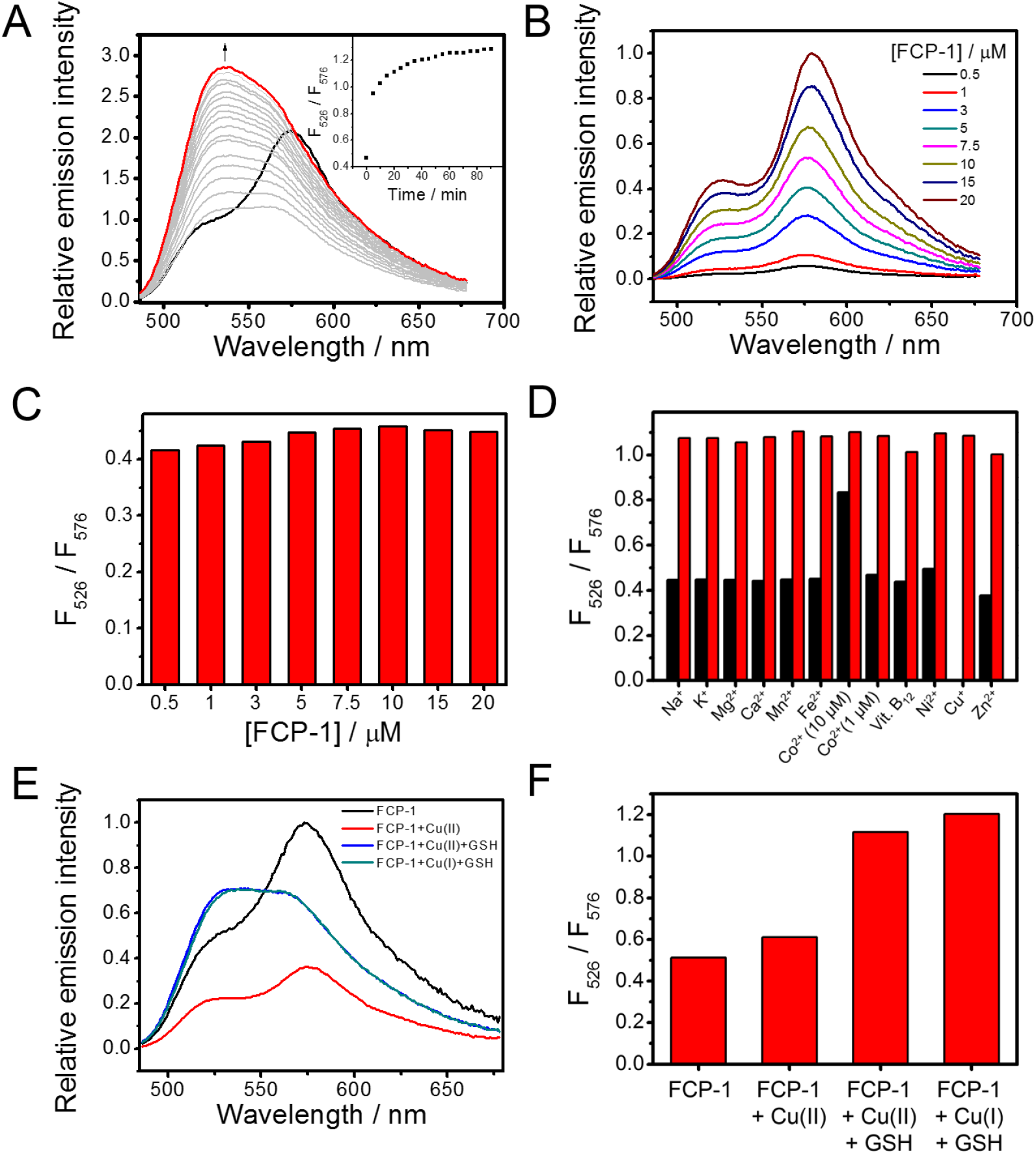
Photophysical properties and Cu(I) selectivity of FCP-1 in aqueous buffer solutions. (A) Changes in corrected emission spectra of FCP-1 (5 μM) in aqueous buffer solution. Inset shows changes in ratiometric emission of FCP-1, *F*_526_ / *F*_576_, with Cu(I) (10 µM) over time. (B) Corrected emission spectra of FCP-1 measured at various concentrations. (C) No significant changes in the *F*_526_ / *F*_576_ emission ratio with varying concentrations of FCP-1 are observed. (D) Changes in *F*_526_ / *F*_576_ ratio of FCP-1 (5 μM) towards various biologically relevant metal ions, as well as cobalt(II)-containing vitamin B12. Black bars represent the ratiometric emission response observed upon addition of competing metal ions (1 mM for Na^+^, K^+^, Mg^2+^ and Ca^2+^, and 10 μM for all the other d-block metal ions as well vitamin B12) after 15 min. Red bars represent the ratiometric emission response upon subsequent addition of 10 μM of Cu(I) after 15 min. (E) Emission spectra and (F) *F*_526_ / *F*_576_ emission ratio of FCP-1 (5 μM) with addition of Cu(II) (10 μM), Cu(II) (10 μM) + GSH (2 mM), or Cu(I) (10 μM) + GSH (2 mM) after 15 min. All spectra were acquired in aqueous buffer (PBS with 2 mM GSH and 40 vol% PEG-400) with λ_ex_ = 458 nm.

Upon addition of Cu(I), FCP-1 showed an increase in fluorescein emission at 526 nm over time, with a concomitant drop in rhodamine fluorescence at 576 nm (Fig. 1A). LC-MS analysis confirms formation of fluorescein methyl ester (**P1**) and rhodamine-TPA (**P2**) fragments upon addition of Cu(I) to a solution of FCP-1 (Fig. S5) and aerobic conditions are required for triggering emission spectral changes (Fig. S6), consistent with a Cu(I)-mediated oxidative cleavage of the benzyl ether C–O bond on the TPA linker of FCP-1. This C-O cleavage leads to separation of the fluorescein donor from rhodamine acceptor and hence results in a decrease in intramolecular FRET efficiency with a ratiometric emission spectral change (Fig. 1A). The ratiometric fluorescence change of FCP-1 at 526 and 576 nm (*F*_526_ / *F*_576_) was rapid and reached saturation after ca. 30 min in the presence of a two-fold excess of Cu(I) (10 μM). Notably, the *F*_526_/ *F*_576_ ratio of FCP-1 showed a linear dose response with Cu(I) addition from 0.01 to 1 µM (Fig. S7A), highlighting the capability of FCP-1 to detect Cu(I) levels in this range by a ratiometric fluorescence change. FCP-1 showed high sensitivity to Cu(I), as revealed by the detectable emission spectral changes at 10 nM of Cu(I) (Fig. S7B) and a detection limit of 16.7 nM based on the 3σ method from 5 independent experiments (64, 65). More interestingly, the basal *F*_526_ / *F*_576_ FCP-1 emission ratio did not show significant changes with varying concentrations of probe (0.5 to 20 µM, Figs. 1B and 1C). This internal self-calibration is advantageous, as FCP-1 was found to be less fluorescent at higher concentrations, presumably due to self-quenching, which is a common issue found in fluorophores (Fig. S8) (66). Indeed, the negligible changes in ratiometric emission of FCP-1 indicated that FRET occurred primarily through an intramolecular mechanism, and concentration-independent ratiometric fluorescence of FCP-1 further highlights its potential in comparative imaging of labile Cu(I) pools in biological specimens by avoiding potential complications arising from variations in probe uptake, sample thickness, and light intensity.

The ratiometric fluorescence change of FCP-1 was selective toward Cu(I) over other biologically relevant metal ions, except free Co(II) at 10 μM, which altered FCP-1 fluorescence modestly (Fig. 1D). Notably, this concentration of exogenous Co(II) is far higher than the physiologically relevant levels of labile Co(II), as this metal ion is known to be tightly bound by proteins. Indeed, FCP-1 showed no significant changes in ratiometric signal in the presence of 10 μM cobalamin (vitamin B_12_) (67), which is the predominant form of Co(II) in mammalian systems, suggesting that interference in the detection of Cu(I) by FCP-1 from Co(II) in biological specimens should be negligible. Competitive experiments further showed the ability of FCP-1 to detect Cu(I) in solutions containing other biologically relevant metal ions (Fig. 1D). In addition to exhibiting metal selectivity, the ratiometric response of FCP-1 is oxidation state-specific for Cu(I), as the probe does not respond ratiometrically to Cu(II) (Figs. 1E and 1F). In further support of this oxidation state selectivity, co-incubation of FCP-1 with Cu(II) and excess glutathione, which readily reduces Cu(II) to Cu(I), resulted in emission spectra and a *F*_526_ / *F*_576_ ratio that is identical to that obtained by treating FCP-1 with Cu(I) and excess glutathione (Figs. 1E and 1F). This oxidation state-specificity is important for the applications of FCP-1 in imaging dynamic changes in intracellular labile Cu(I) pools under redox stress and oncogenic transformation, which we will describe later.

Finally, FCP-1 does not respond to Ag(I), which is a close analog of Cu(I) but lacks the potent oxygen-dependent redox activity (Fig. S9). In view of the low solubility of Ag(I) in the presence of chloride and other halide ions, NaH_2_PO_4_ buffer solution (50 mM, pH 7.4) was used instead of PBS solution in these sets of experiments. Indeed, no significant change in *F*_526_ / *F*_576_ ratio was observed in the presence of Ag(I), while Cu(I) induced a patent increase in *F*_526_ / *F*_576_ ratio of FCP-1 in the same buffer solution (Fig. S9). These data suggest that the observed ratiometric fluorescence changes of FCP-1 upon addition of Cu(I) are attributable to the redox activity of Cu(I). In addition, the presence of reducing agents was found to be crucial for triggering the Cu(I)-mediated ratiometric fluorescence changes of FCP-1 (Fig. S10), and indeed similar fluorescence changes were also observed using ascorbate instead of 2 mM GSH as a reductant (Fig. S11). Together with the observed increase in GSSG (the oxidized form of GSH) levels with a concomitant drop in GSH (the reduced form) levels upon addition of Cu(I) to the solution mixture of FCP-1 and GSH (Fig. S5A), these data support a role for reducing agents in “recycling” the redox-active Cu(I) species for binding to the TPA receptor and triggering subsequent activity-based cleavage of the benzyl ether C–O bond on the linker. This bond cleavage occurs primarily through an oxidative mechanism, because no significant emission spectral changes are observed with Cu(I) addition under anaerobic conditions (Fig. S6). Generation of the fluorescein methyl ester and rhodamine-TPA fragments (Fig. S5) results in a decrease in intramolecular FRET efficiency and an increase in *F*_526_ / *F*_576_ ratio.

### FCP-1 Can Image Labile Cu(I) Pools in Living Cells

Given the metal and oxidation state-selective and sensitive response of FCP-1 to Cu(I), we sought to apply this probe for ratiometric fluorescence imaging of labile Cu(I) pools in living cells. To this end, HEK 293T cells were exposed to varying doses of CuCl_2_ (25-300 µM), washed thoroughly to remove excess copper from the medium, and then incubated with FCP-1 (5 μM) for 45 min and imaged. The copper-supplemented cells showed a statistically significant, dose-dependent increase in *F*_green_ / *F*_orange_ ratio over vehicle control cells (Fig. 2B). In contrast, copper-deficient HEK 293T cells obtained by treatment with a membrane-impermeable copper chelator, bathocuproine sulfonate (BCS; 100 μM), washed, and subsequently incubated with FCP-1 (5 μM) exhibited a statistically significant decrease in *F*_green_ / *F*_orange_ ratio when compared to control cells (Figs. 2C and 2D). The observed increases and decreases in labile Cu(I) levels upon copper supplementation (CuCl_2_) and copper depletion (BCS) treatments, respectively, as measured by FCP-1 imaging were supported by complementary ICP-MS measurements of total copper levels showing expected increases and decreases in the total copper pool (Fig. 2E). Control experiments showed that the cells remained viable as indicated by nuclear staining (Figs. S12 and S13). These data establish that FCP-1 is capable of visualizing both increases and decreases in labile Cu(I) pools in living cells by a ratiometric fluorescence readout.

**Fig. 2.**
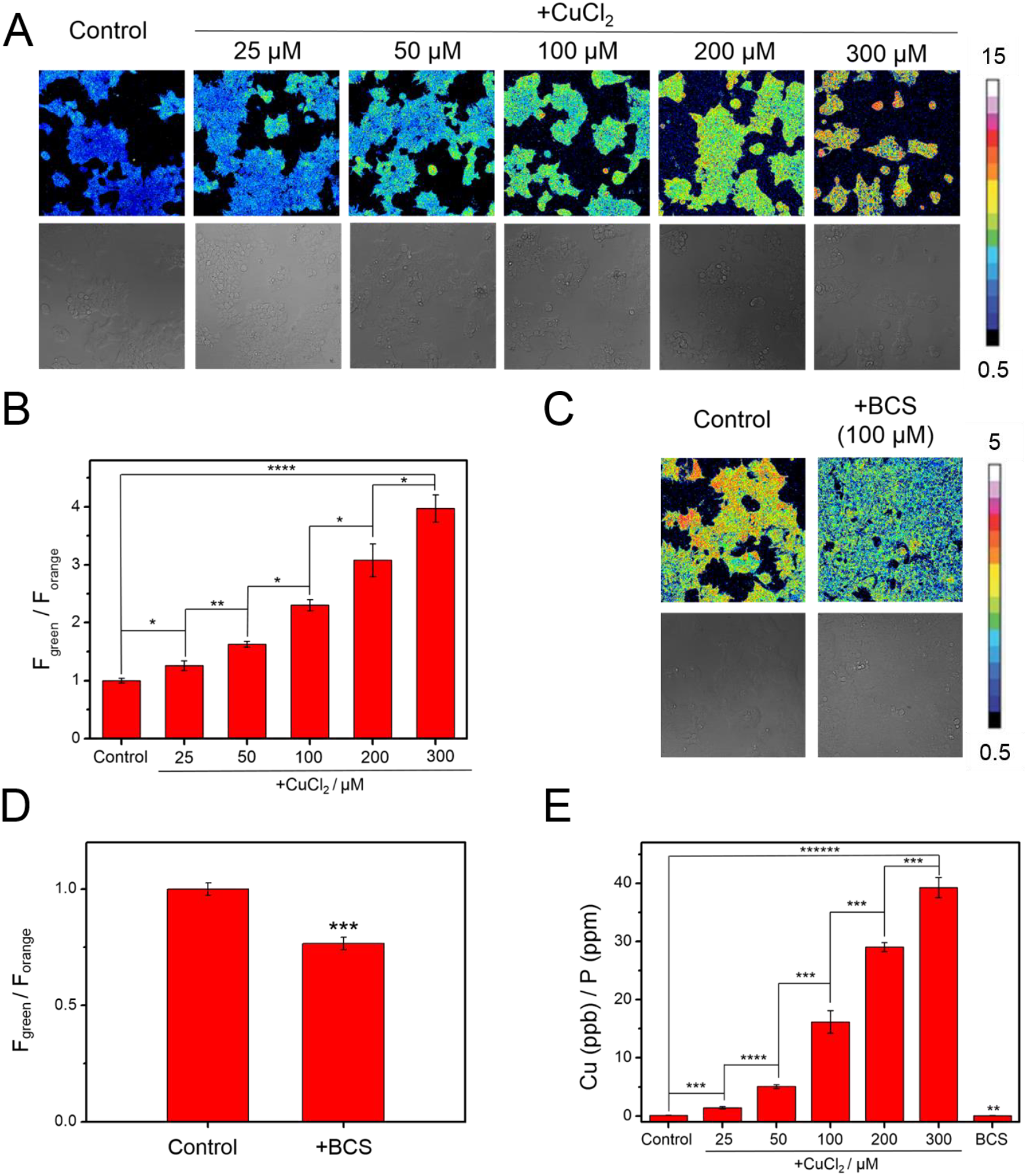
Ratiometric fluorescence imaging of labile Cu(I) levels in living cells using FCP-1. (A) Confocal fluorescence microscopy images of HEK 293T cells pretreated with solvent vehicle control or varying concentrations of CuCl_2_ in complete medium for 18 h. The cells were then washed with PBS, incubated with FCP-1 (5 μM) in DPBS for 45 min and then imaged. (B) Average cellular ratiometric emission ratios, represented by *F*_green_/*F*_orange_, as determined from imaging experiments in (A) performed in triplicate. (C) Confocal fluorescence microscopy images of HEK 293T cells pretreated with solvent vehicle control or BCS (100 μM) in complete medium for 18 h. The cells were then washed with PBS, incubated with FCP-1 (5 μM) in DPBS for 45 min and then imaged. (D) Average cellular ratiometric emission ratios of FCP-1, *F*_green_/*F*_orange_, as determined from experiments in (C) performed in triplicate. Panels (A) and (C) are displayed in pseudocolor and referenced to the basal control for that experiment and independently from each other. λ_ex_ = 458 nm. (E) ICP-MS measurement to determine total cellular ^63^Cu levels in HEK293T cells under copper supplementation and depletion relative to the control (with normalization of different cell numbers by total cellular ^31^P level). Error bars denote standard derivation (SD; *n* = 3). \**p* < 0.05, \*\**p* < 0.01, \*\*\**p* < 0.001, \*\*\*\**p* < 0.0001 and \*\*\*\*\**p* < 0.00001.

### FCP-1 Can Monitor Differences in Labile Cu(I) Levels in a Genetic Cell Model of Copper Misregulation

Having established that FCP-1 is sensitive to endogenous pools of labile Cu(I) under basal conditions and can respond to changes in labile Cu(I) pools upon pharmacological copper supplementation and/or depletion, we sought to explore the application of FCP-1 to visualize alterations in endogenous labile Cu(I) pools through genetic manipulation. To this end, we performed comparative FCP-1 imaging in mouse embryonic fibroblast cells that are either Cu-replete (*Ctr1*^+/+^ MEFs) or Cu-deficient (*Ctr1*^−/−^ MEFs) through knockout of the high-affinity copper transporter CTR1 (68–70). As expected, *Ctr1*^+/+^ MEFs incubated with FCP-1 exhibited higher *F*_green_ / *F*_orange_ ratios as compared to the *Ctr1*^−/−^ MEFs (*p* < 0.01; Figs. 3A and 3B), indicating that FCP-1 can precisely detect a relative decrease in endogenous levels of labile Cu(I) in cells deficient in copper uptake.

**Fig. 3.**
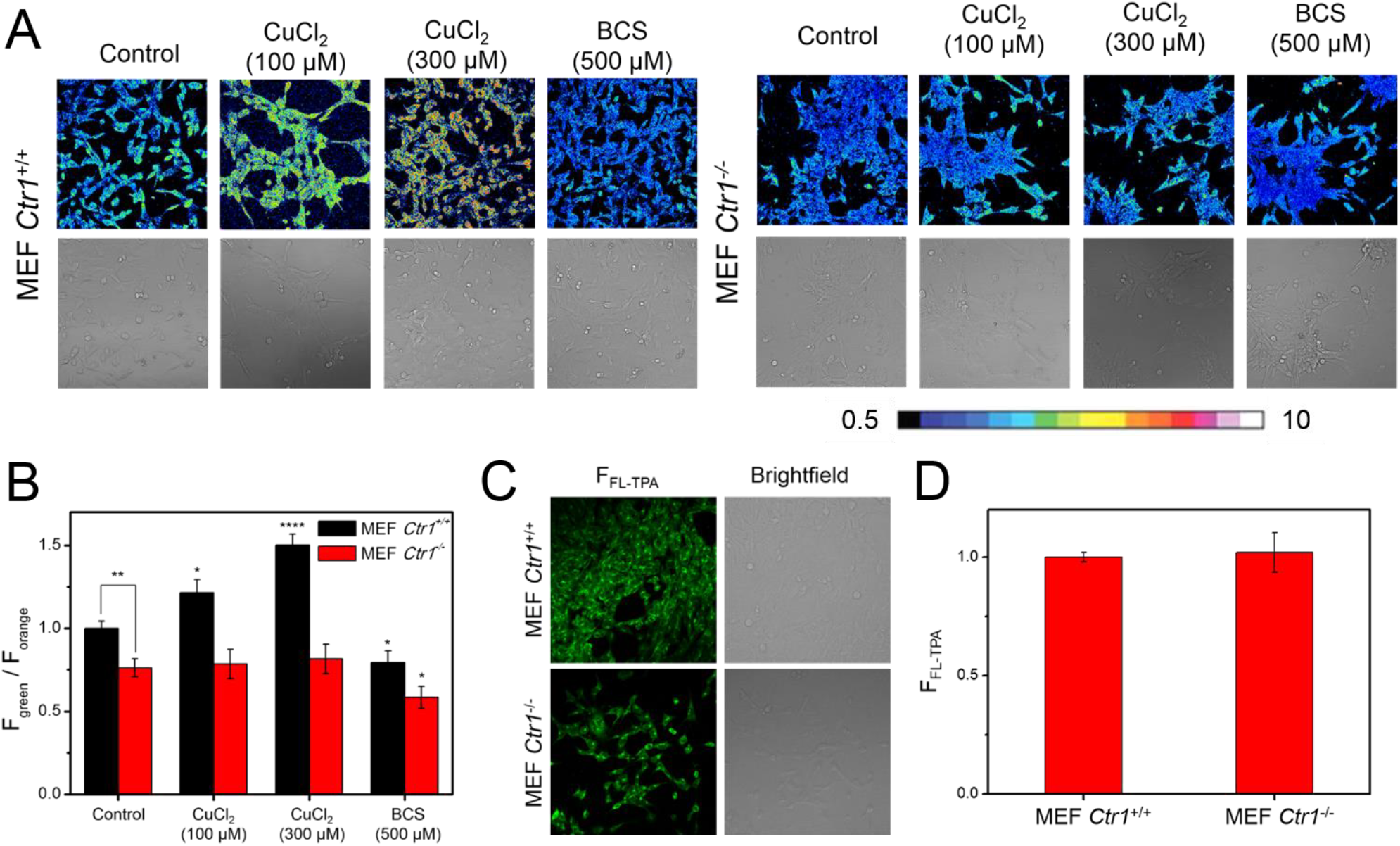
FCP-1 Enables Comparative Imaging of Labile Cu(I) Levels in Live cells With Altered Expression Levels of the High-Affinity Copper Uptake Protein CTR1. (A) Confocal fluorescence microscopy images of *Ctr1*^+/+^ MEFs and *Ctr1–/–* MEFs treated with solvent control, CuCl_2_ (100 or 300 μM) or BCS (500 μM) in complete medium for 8 h, washed with complete medium and PBS, incubated with FCP-1 (5 μM) in DPBS for 45 min, and then imaged with λ_ex_ = 458 nm. (B) Average cellular ratiometric emission ratios of FCP-1, *F*_green_/*F*_orange_, as determined from experiments performed in triplicate. (C) Confocal fluorescence microscopy images of *Ctr1*^+/+^ MEFs and *Ctr1*^−/−^ MEFs incubated with FL-TPA (5 μM) in DPBS for 45 min. The cells were then imaged with λ_ex_ = 488 nm. (D) Average cellular fluorescence intensity of FL-TPA, *F*_FL-TPA_, determined from experiments performed in triplicate with λ_ex_ = 488 nm. Error bars denote SD (*n* = 3). \**p* < 0.05, \*\**p* < 0.01 and \*\*\**p* < 0.001.

To further validate the capability of FCP-1 to monitor differential levels of labile Cu(I) in cells, *Ctr1^+/+^* MEFs incubated with varying concentrations of CuCl_2_ exhibited a dose-dependent increase in the FCP-1 *F*_green_ / *F*_orange_ ratio. In contrast, no statistically significant changes in *F*_green_ / *F*_orange_ ratios were observed in *Ctr1*^−/−^ MEFs incubated with CuCl_2_ and then visualized with FCP-1 (Figs. 3A and 3B). In meantime, the cellular viability was maintained, as confirmed with the nuclear stain DRAQ5 (Figs. S14 and S15). The difference of FCP-1 ratiometric fluorescence response to exogenous copper supplementation between the wildtype and *Ctr1* knockout MEFs is consistent with inefficient copper uptake rather than changes in copper efflux, as treatment of *Ctr1*^−/−^ MEFs with the membrane-impermeable copper chelator BCS resulted in a reduction in *F*_green_ / *F*_orange_ ratio, similar to what is observed in *Ctr1*^+/+^ MEFs (Figs. 3A and 3B). Complementary ICP-MS measurements confirm that total copper levels in *Ctr1*^−/−^ MEFs are lower than *Ctr1*^+/+^ MEFs (Fig. S16). Importantly, the ratiometric response of FCP-1 is key to its ability to distinguish between basal and decreased levels of labile Cu(I). Indeed, the control probe FL-TPA, which exhibits a “turn-on” fluorescence response without ratiometric self-calibration, was unable to distinguish differences in basal labile Cu(I) levels in *Ctr1*^+/+^ MEFs and *Ctr1*^−/−^ MEFs (Figs. 3C and 3D), while being able to detect changes in labile copper with exogenous copper addition (Fig. S17). This result can be explained by potential differences in probe loading into the two MEF cell lines, further highlighting the self-calibrating nature of the ratiometric probe FCP-1 as a critical feature for accurate detection of analyte levels across different, variable biological specimens. Finally, we tested whether FCP-1 could disrupt labile Cu pools by measuring CCS protein levels in MEFs treated with FCP-1 or BCS, the latter as a positive control to reduce cellular Cu content and promote CCS protein stability. Indeed, CCS protein levels were unaltered in cells treated with a concentration of FCP-1 sufficient to visualize cellular labile Cu pools, while BCS treatment increased CCS, as expected (Fig. S18), confirming that FCP-1, itself, does not alter cellular Cu levels under these conditions. The data demonstrate that FCP-1 can monitor basal levels of endogenous labile Cu(I) and depletion of such pools in response to reduced cellular copper uptake through either genetic or pharmacologic means.

### FCP-1 Identifies a Dynamic Change in Labile Cu(I) Versus Total Copper Pools in Living Cells Under Redox Stress

In view of the oxidation state-specificity of FCP-1 toward Cu(I), we investigated the impact of cellular redox status on labile Cu(I) pools, which remains insufficiently understood. First, we validated the ability of FCP-1 to detect changes in labile Cu(I) levels in HeLa cells with CuCl_2_ and BCS treatments (Fig. 4A). We then monitored the effect of addition of ascorbate, which is a powerful reducing agent, on intracellular labile Cu(I) pools. Interestingly, we observed a statistically significant increase in FCP-1 *F*_green_ / *F*_orange_ ratio, suggesting an increase in labile Cu(I) levels in the ascorbate-treated cells compared to the control cells (Fig. 4B) (31). Since the accompanying ICP-MS measurements showed no changes in total copper levels in cells treated with ascorbate compared to vehicle controls (Fig. 4C), the increase in *F*_green_ / *F*_orange_ can be ascribed to an elevation in labile Cu(I) levels. Several potential factors could contribute to this observation in response to this exogenous reducing agent, including a shift in the global or local Cu(I)/Cu(II) redox balance, a lowered cellular or subcellular capacity to buffer Cu(I), and/or relocalization/recomparmentalization of labile Cu(I) pools.

**Fig. 4.**
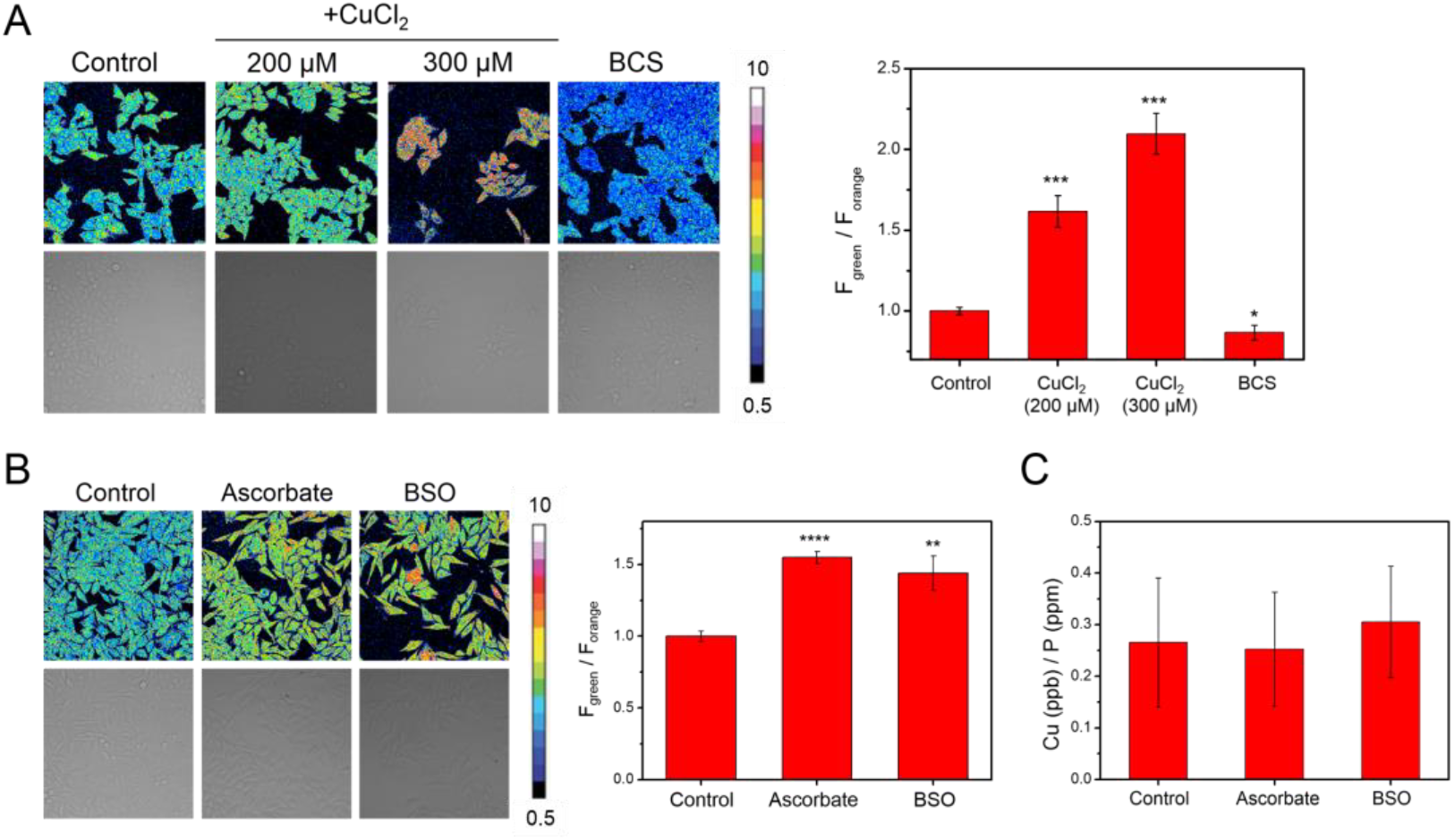
FCP-1 Identifies Changes in Labile Cu(I) But Not Total Copper Levels in Live Cells Under Redox Stress. Confocal fluorescence microscopy images of HeLa cells pretreated with (A) solvent control, CuCl_2_ (200 or 300 μM), or BCS (100 μM) in complete medium for 18 h, or (B) solvent control, ascorbate (1 mM), or BSO (1 mM) in complete medium for 4 h. The cells were then washed with PBS, incubated with FCP-1 (5 μM) in DPBS for 45 min and imaged with λ_ex_ = 458 nm. The bar charts show average cellular ratiometric FCP-1 emission, *F*_green_/*F*_orange_, determined from experiments performed in triplicate. (C) Total cellular ^63^Cu levels were determined by ICP-MS experiments (with normalization of different cell numbers by total cellular ^31^P level). Error bars denote SD (*n* = 3). \**p* < 0.05, \*\**p* < 0.01 and \*\*\**p* < 0.001 and show no change with either ascorbate or BSO treatment.

With these results in hand, we next sought to utilize FCP-1 to explore potential connections between the maintenance of labile Cu(I) pools and endogenous central redox mediators in the cell. In this context, glutathione (GSH) is a ubiquitous, naturally-occurring antioxidant and a key small molecule for maintaining proper cellular redox status (71). The rate-determining step of GSH biosynthesis involves glutamate cysteine ligase (GCL), which can be inhibited by buthionine sulfoximine (BSO) (72). Interestingly, we observed that the FCP-1 *F*_green_ / *F*_orange_ ratio was higher in BSO-treated HeLa cells when compared to control cells (Fig. 4B), indicating an increase in labile Cu(I) levels when GSH synthesis is blocked. Moreover, total intracellular copper levels did not differ significantly between the BSO-pretreated cells and control cells as shown by ICP-MS (Fig. 4C), showing that the dynamic changes are in the labile Cu(I) pool rather than an overall increase or decrease in the total copper pool. On balance, we caution against overinterpreting these data as supporting a simple and direct Cu(I)-glutathione complexation model, as although aqueous solutions of FCP-1 and Cu(I) were found to show higher *F*_526_ / *F*_576_ ratios at lower GSH concentrations *in vitro* (Fig. S19), the variable *K*_d_ values and copper-ligand stoichiometries obtained under different buffer conditions do not rule out more sophisticated interplay between metal and redox homeostasis pathways (7, 22, 73). Indeed, GSH is not just a reducing agents but also has been reported to interact with metallochaperones like ATOX1 or metal-buffering/scavenging proteins such as metallothionein to form complexes that can bind copper tightly (73–76). Therefore, the observed increase in labile Cu(I) levels resulting from GSH deficiency may be operating through more complex, indirect mechanisms, an area that warrants further study.

To further investigate correlations between labile Cu(I) pools and altered GSH metabolism, we prepared two MEF cell lines for study, where one line exhibits lower levels of total GSH via stable genetic knockdown of endogenous *Gclc* (the catalytic subunit of the rate limiting enzyme GCL in GSH synthesis) and the other line exhibits lower levels of GSH and a lower ratio of reduced to oxidized GSH (GSH/GSSG) via knockdown of glutathione reductase (*Gsr*; the NADPH dependent-enzyme that cleaves the disulfide bond on two oxidized GSH molecules) (71, 77). As expected, shRNA-mediated knockdown of endogenous *Gclc* (Figs. 5A and B) significantly reduced total GSH levels to a similar degree to MEFs treated with BSO, which potently inhibits GCLC enzymatic function, when compared to non-targeting scramble control (*Scr*) MEFs (Fig. 5C). Importantly, no significant change in the GSH/GSSG ratio is observed in these *Gclc* knockdown cells despite the lower total GSH levels (Fig. 5D). In contrast, genetic knockdown of *Gsr* via shRNA significantly lowered both total GSH levels and the GSH/GSSG ratio, while treatment with the GSR inhibitor, Carmustine (bis-chloroethylnitrosourea, BCNU) only significantly lowered the GSH/GSSH ratio (Figs. 5C, 5D, 5E and 5F), indicating fluctuations in intracellular redox balance to a more oxidizing environment in the absence of the ability to regenerate GSH from GSSG.

**Fig. 5.**
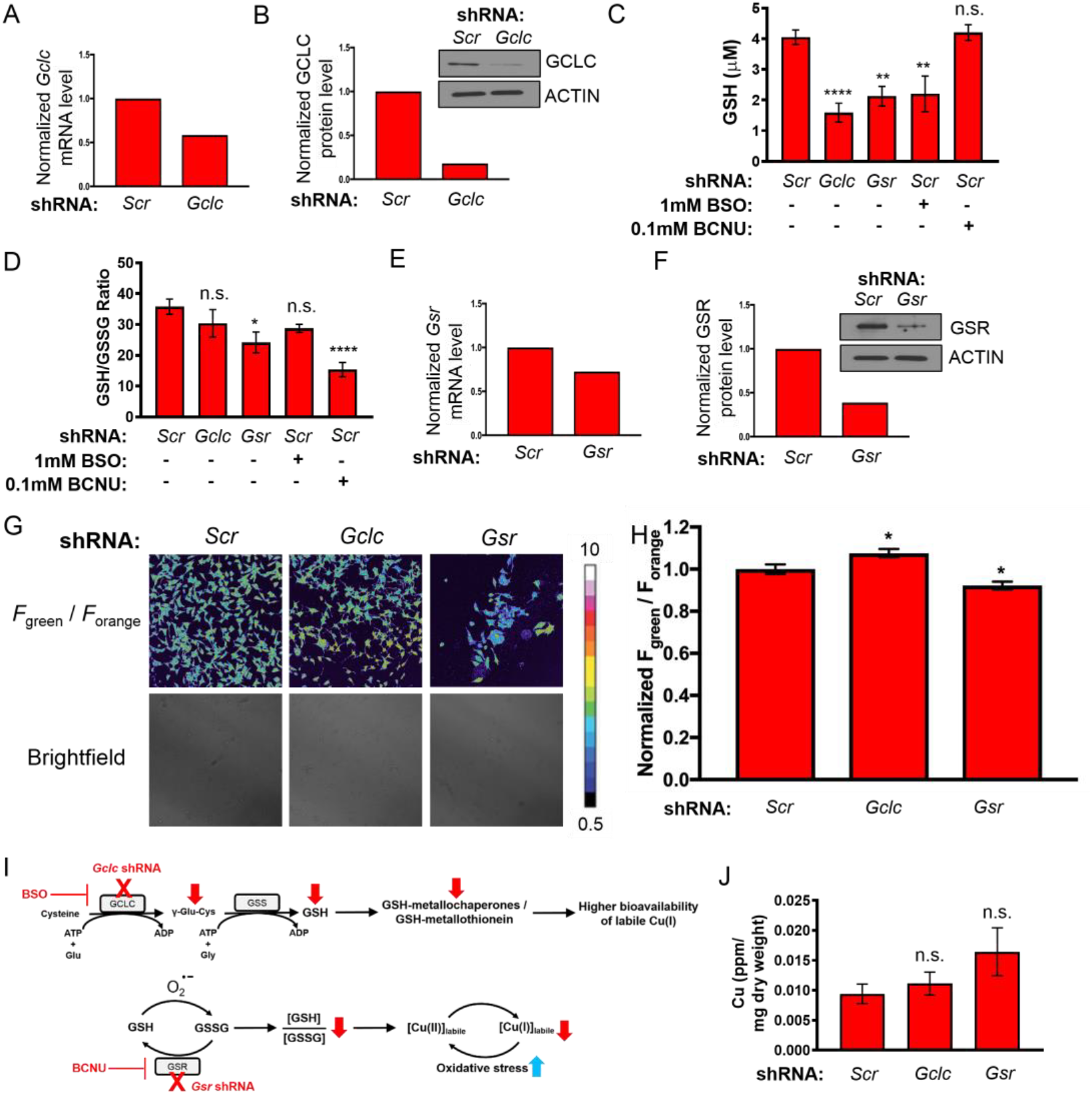
FCP-1 Imaging Reveals Elevation or Depletion of Labile Cu(I) Pools Induced by Genetic Alterations in Total Glutathione (GSH) or Reduced/Oxidized Glutathione Ratio (GSH/GSSG). (A) Normalized quantitative PCR (qPCR) expression of *Gclc* mRNA and (B) immunoblot detection of GCLC or ACTIN in MEFs stably expressing non-targeting control shRNA (*Scr*) and *Gclc* shRNA. (C) and (D) display quantification of total cellular glutathione (GSH, μM) or the ratio of GSH to GSSG (GSH/GSSG) ± s.e.m. from MEFs stably expressing *Scr*, *Gclc* shRNA, *Gsr* shRNA, or *Scr* treated with 1 mM BSO or 0.1 mM BCNU. The GSH and GSH/GSSG levels were determined using GSH/GSSG-Glo™ Assay kit, and the results were compared using one-way ANOVA followed by a Dunnett’s multiple comparison test (n ≥ 11). (E) qPCR expression of *Gsr* mRNA and (F) immunoblot detection of GSR or ACTIN in MEFs stably expressing *Scr* and *Gs* shRNA. (G) Representative confocal fluorescence images of MEFs stably expressing *Scr*, *Gclc* shRNA, or *Gsr* shRNA incubated by FCP-1. (H) Quantification of normalized mean FCP-1 fluorescence emission (*F*_green_/*F*_orange_) ± s.e.m. Data are from analysis of 98 individual cells. (n=98) (I) Proposed model for alterations in glutathione biosynthesis enzymes and redox stress on the bioavailability of labile Cu(I) pools. (J) Total Cu levels (parts per million, ppm) detected by ICP-MS from MEFs stably expressing *Scr*, *Gclc* shRNA, or *Gsr* shRNA per sample weight ± s.e.m (n = 6). Results were compared using a one-way ANOVA followed by a Dunnett’s multi-comparisons test. \**p* < 0.05, \*\**p* < 0.01 and \*\*\*\**p* < 0.0001; n.s. = not statistically significant.

After validating the successful manipulation of either GSH levels or the GSH/GSSG ratio, FCP-1 was applied to probe the relative levels of labile Cu(I) across this panel of genetically defined cell lines. *Gclc* knockdown cells showed a higher FCP-1 *F*_green_ / *F*_orange_ ratio as compared to the *Scr* cells, while a lower FCP-1 *F*_green_ / *F*_orange_ ratio was observed in *Gsr* knockdown cells as compared to the control (Figs.5G and 5H). Since *Gclc* knockdown inhibits the enzyme involved in the rate-determining step of GSH biosynthesis and mimics pharmacological treatment with BSO (Fig. 4B), we attribute the observed increase in *F*_green_ / *F*_orange_ ratio to the decrease in total GSH levels. The minimal change in GSH/GSSG ratio in this *Gclc* knockdown cell line suggests that the overall cellular redox state is unaffected and instead points to decreases in levels of labile Cu(I)-buffering systems (Fig. 5I). Included are potential GSH-metallochaperone or GSH-metallothionein adducts, both of which can influence loosely-bound vs tightly-bound Cu(I) pools. On the other hand, *Gsr* knockdown cells showed a lower GSH/GSSG ratio (Fig. 5D), which reflects a more overall oxidizing intracellular environment and may lead to a shift in the dynamic balance of Cu(I/II) pools, which results in a decrease in labile Cu(I) levels (Fig. 5I). Although total GSH levels also decrease in *Gsr* knockdown cells, which may increase bioavailability of labile Cu(I) to some extent, the statistically significant decrease in the FCP-1 *F*_green_ / *F*_orange_ ratio relative to control indicates an overall labile Cu(I) deficiency and suggests that the redox contribution of more GSSG relative to GSH is the dominant contributor. Because the FCP-1 activity-based sensing mechanism relies on an oxygen-dependent redox reaction with Cu(I), we performed an additional set of control experiments using a previously reported copper fluorescent turn-on probe, Copper Fluor-4 (CF4), that operates by a direct, reversible Cu(I) binding-based mechanism (14). CF4 showed increased fluorescence in *Gclc* knockout cell lines but decreased fluorescence in *Gsr* knockout cells, which agrees well with those obtained with FCP-1 (Fig. S20), providing further support for changes in labile Cu(I) pools induced by alterations in GSH and GSH/GSSG status. Finally, whereas the labile Cu(I) pools change in MEFs lacking *Gclc* or *Gsr*, no changes in total copper content were observed as measured by ICP-MS (Fig. 5J). These experiments reveal that labile Cu(I) pools can be elevated and/or depleted in a glutathione-dependent manner depending on both total GSH levels as well as GSH/GSSG redox ratios.

### Oncogenic BRAF^V600E^ and KRAS^G12^ Mutations Lead to Labile Cu(I) Deficiency Due in Part to Decreased Expression of GSR and Diminished Ability to Regenerate GSH

Noting the growing connections between copper and kinase signaling pathways involved in oncogenesis, we next sought to apply the identification of copper-glutathione crosstalk to tumorigenesis. Indeed, rapid proliferation associated with tumorigenesis requires cancer cells to increase energy metabolism and initiate cellular mechanisms to respond to oncogenic stress to sustain unrestricted tumor growth, survival, and therapeutic resistance. For example, many solid tumors display increased reactive oxygen species (ROS) production in response to higher energic needs and mitochondrial dysfunction. As a result, this induces an antioxidant program to detoxify from ROS and maintain a more robust redox balance (78). For example, MEFs expressing conditional oncogenic alleles of KRAS^G12D^ or BRAF^V619E^ display reduced levels of intracellular ROS with concomitant increases in mRNA expression of the glutathione metabolism genes *Gclc* and *Gclm* (79). Moreover, ectopic expression of KRAS^G12D^ increased total GSH levels yet a lower GSH/GSSG ratio. However, whether oncogenic mutations in *BRAF* or *KRAS* alter the intracellular pools of labile Cu(I) by disrupting intracellular redox balance, particularly through glutathione pathways, remains insufficiently understood.

To address this question, immortalized MEFs were transformed with BRAF^V600E^ or KRAS^G12D^ and levels of intracellular labile Cu(I) were assessed with ratiometric FCP-1 imaging. Interestingly, we observed a decrease in the average cellular ratiometric fluorescence from both BRAF^V600E^ and KRAS^G12D^-transformed MEFs cells as read by FCP-1 *F*_green_ / *F*_orange_ ratio when compared to control congeners (Figs. 6A and 6B). These findings were phenocopied in live-cell imaging experiments using the complementary CF4 probe, suggesting that loosely-bound pools of intracellular Cu(I) are diminished upon oncogenic transformation (Figs. S21A and S21B). In agreement, we found that CCS protein stability, which is degraded in a Cu-dependent fashion, is increased in cells stably expressing these oncogenes or when cells are treated with the Cu chelator BCS, as a positive control, suggesting a depletion of labile intracellular Cu pools (Fig. 6C). To determine whether the observed labile Cu(I) deficiency as detected by live-cell FCP-1 and CF4 imaging was reflected in changes in total cellular Cu levels, ICP-MS was performed on control, BRAF^V600E^, or KRAS^G12D^ expressing MEFs, and these cells showed unchanged total Cu levels (Fig. 6D), confirming that labile Cu(I) pool is selectively altered rather than the total Cu pool. Furthermore, *Ctr1* transcript levels in both BRAF^V600E^ and KRAS^G12D^ cells remain similar to control cells, suggesting that the observed decrease in labile Cu(I) levels is not due to changes in cellular Cu import (Fig. 6E). Although *Atp7a* transcript levels were insignificantly lower in both BRAF^V600E^ and KRAS^G12D^ cells (Fig. 6F), these changes in *Atp7a* mRNA expression may reflect the cells sensing lower bioavailable Cu(I) levels. As an alternative hypothesis to differential expression of the Cu transporters CTR1 and ATP7A to decrease labile Cu(I) levels, recent studies have established that decreasing the levels of CTR1 or introducing surface accessible mutations in MEK1, which functions downstream of oncogenic BRAF^V600E^, as an approach to disrupt Cu binding decreased BRAF^V600E^-driven signaling and tumor growth (5, 80). Thus, the Cu-MEK1/2 interaction is essential for BRAF^V600E^-driven signaling and tumorigenesis and decreased labile Cu levels in BRAF^V600E^-transformed MEFs could potentially be driven by increased delivery of Cu to MEK1/2 to support hyperactivated MAPK signaling. To experimentally address this possibility, BRAF^V600E^-transformed MEFs were engineered to stably express scramble shRNA or *Mek1* shRNA with RNA-mediated interference-resistant wild-type MEK1 or a Cu-binding mutant of MEK1 (MEK1^CBM^) and the ratiometric fluorescence from FCP-1 in these cell lines was compared to non-transformed MEFs. While BRAF ^V600E^ expression reduced *F*_green_ / *F*_orange_ by ~20%, the decrease was maintained in the presence of both wild-type MEK1 and MEK1^CBM^, suggesting that the observed decrease in the labile Cu(I) pool is independent of MEK1/2 Cu binding (Fig. S22).

**Fig. 6.**
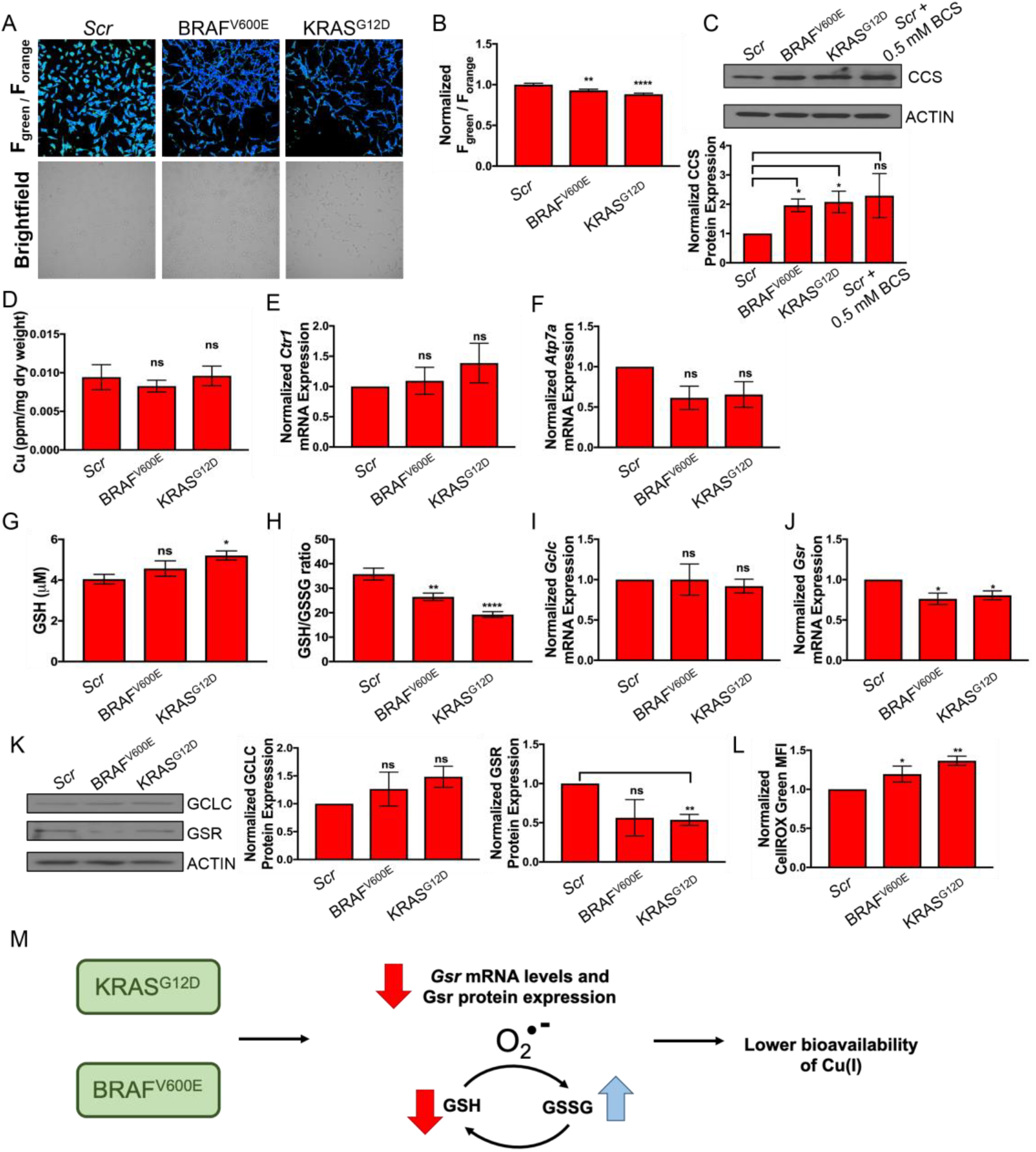
Oncogenic BRAF^V600E^ and KRAS^G12D^ Lead to Decreases in Labile Cu(I) Pools by Decreasing Levels of *Gsr* and Increasing Expression of CCS. (A) Representative ratiometric live-cell imaging of FCP-1 on MEFs stably expressing non-targeting control shRNA (*Scr*), *BRAF^V600E^* cDNA, or *KRAS^G12D^* cDNA. (B) Quantification of mean FCP-1 ^fluorescence emission (*F*^green^/*F*^orange^) ± s.e.m. from MEFs stably^ expressing *Scr*, *BRAF^V600E^* cDNA, or *KRAS^G12D^* cDNA. Results were compared using a one-way ANOVA followed by a Dunnett’s multi-comparisons test. Data are from analysis of 105 or more individual cells (n≥105). (C) Immunoblot detection of CCS or ACTIN in MEFs stably expressing *Scr*, *BRAF^V600E^* cDNA, or *KRAS^G12D^* cDNA or *Scr* treated with 0.5 mM BCS. Quantification: ΔCCS/ACTIN normalized to *Scr*. Results from three independent biological replicates were compared using an unpaired t-test. (D) Quantification of mean Cu levels (parts per million, ppm) detected by ICP-MS from MEFs stably expressing *Scr*, *BRAF^V600E^* cDNA, or *KRAS^G12D^* cDNA per sample weight ± s.e.m. (n=5). Results were compared using a one-way ANOVA followed by a Dunnet’s multi-comparisons test. (E and F) qPCR expression of *Ctr1* or *Atp7a* mRNA from MEFs stably expressing *Scr*, *BRAF^V600E^* cDNA, or *KRAS^G12D^* cDNA. (n=4). Results were compared using a one-way ANOVA followed by a Dunnet’s multi-comparisons test. (G and H) Quantification of total cellular glutathione (GSH, μM) or the ratio of GSH to GSSG (GSH/GSSG) ± s.e.m. from MEFs stably expressing *Scr*, *BRAF^V600E^* cDNA, or *KRAS^G12D^* cDNA. Results were compared using one-way ANOVA followed by a Dunnett’s multiple comparison test (n≥11). Data are from four biological replicates conducted in triplicate. (I and J) qPCR expression of *Gclc* or *Gsr* mRNA from MEFs stably expressing *Scr*, *BRAF^V600E^* cDNA, or *KRAS^G12D^* cDNA. Results were compared using one-way ANOVA followed by a Dunnett’s multiple comparison test (n=4). Data are from four biological replicates conducted in triplicate. (K) Immunoblot detection of GCLC, GSR or ACTIN in MEFs stably expressing *Scr*, *BRAF^V600E^* cDNA, or *KRAS^G12D^* cDNA. Quantification: ΔGCLC/ACTIN or ΔGSR/ACTIN normalized to *Scr*. Results from three independent biological replicates were compared using an unpaired t-test. (L) Flow cytometry analysis of the level of reactive oxygen species quantified by the mean CellROX Green fluorescent intensity ± s.e.m. from MEFs stably expressing *Scr*, *BRAF^V600E^* cDNA, or *KRAS^G12D^* cDNA. Results were compared using a one-way ANOVA followed by a Dunnett’s multiple comparison test (n≥11) \**p* < 0.05, \*\**p* < 0.01, \*\*\**p* < 0.001, and \*\*\*\**p* < 0.0001; ns = not statistically significant. (M) Proposed effects of oncogenic activations of BRAF^V600E^ or KRAS^G12D^ on the bioavailability of labile Cu(I) pools.

Given our collective results that indicate that Cu(I) appears less bioavailable in the context of oncogenic transformation, we hypothesized that differences in glutathione metabolism may be driving the phenotype. To this end, we tested total GSH levels and the GSH/GSSG ratios in MEFs transformed with oncogenic BRAF^V600E^ or KRAS^G12D^ and found that expression of either oncogene increased total GSH levels, although only statistically significant in the KRAS^G12D^ cells (Fig. 6G). To our surprise, GSH/GSSG ratios were significantly lower in the BRAF^V600E^ and KRAS^G12D^ stably expressing MEFs when compared to control cells (Fig. 6H). Since the BRAF^V600E^- and KRAS^G12D^-transformed MEFs phenocopied the genetic knockdown of *Gsr* with respect to reduced ratiometric fluorescence from FCP-1 and a diminished GSH/GSSG ratio, *Gsr* mRNA levels and protein expression were interrogated. When normalized to control cells, *Gsr* transcript levels were significantly reduced upon stable expression of either BRAF^V600E^ or KRAS^G12D^, whereas *Gclc* transcript levels remained relatively unchanged amongst them (Figs. 6I and 6J). Parallel with *Gsr* transcript levels, GSR protein expression was reduced in the BRAF^V600E^ and KRAS^G12D^ cells when compared to control cells (Fig. 6K). In contrast, GCLC protein levels were higher in the BRAF^V600E^ and KRAS^G12D^ cells when compared to control cells (Fig. 6K). To further support the connection between oncogenesis, copper homeostasis, and alterations in glutathione metabolism enzyme expression, both BRAF^V600E^ and KRAS^G12D^ significantly increased the levels of CellROX green fluorescence (Fig. 6L), suggesting that oncogenic transformation increases ROS and in turn reduces the labile Cu(I) pool. From these data, we conclude that BRAF^V600E^ and KRAS^G12D^-transformed cells exhibit a labile Cu(I) deficiency in part due to decreased levels of *Gsr* and increased levels of ROS, which both may foster a more oxidized cellular environment due to the inability to regenerate GSH (Fig. 6M).

## Concluding Remarks

The complex and dynamic interplay between labile and total metal pools has implications for metal homeostasis and metal dysregulation in healthy and disease states. Copper provides a prime example of this signal/stress dichotomy, as its potent redox activity makes it an essential nutrient but this same chemical reactivity makes loss of copper homeostasis a danger to the redox balance of the cell. To address the need for new chemical tools to probe labile copper pools to be used in conjunction with complementary traditional techniques for total copper status, we have developed FCP-1, a first-generation activity-based ratiometric FRET probe that enables selective imaging of labile Cu(I) pools in live cells with a self-calibrating response. FCP-1 utilizes a classic bioinorganic TPA ligand motif for Cu(I)-dependent cleavage to regulate FRET between fluorescein donor and rhodamine acceptor units in a dose-dependent manner, affording high metal and oxidation state specificity for Cu(I). Owing to its ratiometric response, FCP-1 is capable of visualizing dynamic changes in labile Cu(I) levels in live cells with copper supplementation and/or chelation, as well as monitoring decreases in endogenous labile Cu(I) levels in MEF cells deficient in the copper transporter CTR1. Indeed, the ratiometric readout of FCP-1 with internal self-calibration of two emission signatures is the key distinguishing feature to monitor the basal levels of labile Cu(I) across different cell lines, as the control probe FL-TPA that utilizes the same activity-based trigger but responds only in intensity mode at one emission readout cannot discriminate labile Cu(I) status between CTR1 wildtype and CTR knockout lines owing to variability in probe loading.

Against this backdrop, we applied FCP-1 to visualize alterations in labile Cu(I) pools under situations of redox stress, with specific focus on the nexus between copper and glutathione homeostasis. Indeed, we observed elevated levels of labile Cu(I) in *Gclc* knockdown cells, which possess a lower amount of total GSH compared to matched wildtype controls, whereas we detected lower levels of labile Cu(I) in *Gsr* knockdown cells relative to control congeners. These imaging data track with the significant decrease in the GSH/GSSG ratio in *Gsr* knockdown cells but not in *Gclc* knockdown cells. Moreover, we identified a decrease in the GSH/GSSG ratio in MEFs transformed with either oncogenic BRAF^V600E^ or KRAS^G12D^, which in turn induces a labile Cu(I) deficiency. Coupled with the fact that total copper levels remain unchanged in the *Gclc* or *Gsr* knockdown and oncogenic BRAF^V600E^ or KRAS^G12D^ cell lines and that we also observe higher expressions of the CCS protein, which can bind copper tightly and lower labile Cu(I) levels, the ratiometric FCP-1 fluorescence measurements show that the labile Cu(I) pool is selectively affected by oncogene transformation over the total copper pool. By identifying connections between the labile Cu(I) pool, glutathione metabolism, and oncogenic transformation, this work provides a starting point to further interrogate how and why depletion of labile Cu(I) levels can contribute to cancer development and progression. Finally, this study motivates the development of activity-based sensing probes for ratiometric detection of a broader set of target analytes in biological settings where self-calibration can prove advantageous.

## Materials and Methods

Full materials and procedures for the synthesis of compounds, spectroscopic characterization, cellular imaging, generation of cell lines, and cell analysis are described in SI Appendix.

## Author Information

### Author Contributions

C.Y.-S.C., D.C.B. and C.J.C. designed research; C.Y.-S.C., J.M.P., S.L., T.T. and J.M.D. performed research; C.Y.-S.C. and S.L. contributed new reagents, and J.M.P., T.T. and J.M.D. contributed new models; C.Y.-S.C., J.M.P., S.L., T.T., J.M.D., D.C.B. and C.J.C. analyzed data; C.Y.-S.C. and C.J.C. wrote the paper with contributions from all authors. ^1^These authors contributed equally.

## Acknowledgments

We thank National Institutes of Health (NIH) (Grant GM79465 to C.J.C. and Grant GM124749 to D.C.B.) and Pew Charitable Trust Scholars Program in Biomedical Science (Award #50359 to D.C.B.) for support. C.Y.-S.C. acknowledges the Croucher Foundation for a postdoctoral fellowship. J.M.P. acknowledges the American Cancer Society for postdoctoral fellowship (131203PF1714701CCG). T.T. acknowledges the National Cancer Institute (NCI) for predoctoral fellowship (F31CA243294). J.M.D. acknowledges the National Science Foundation for predoctoral fellowship (2017241243). C.J.C. is an Investigator with the Howard Hughes Medical Institute and a CIFAR Senior Fellow.

---

**Scheme 1.**
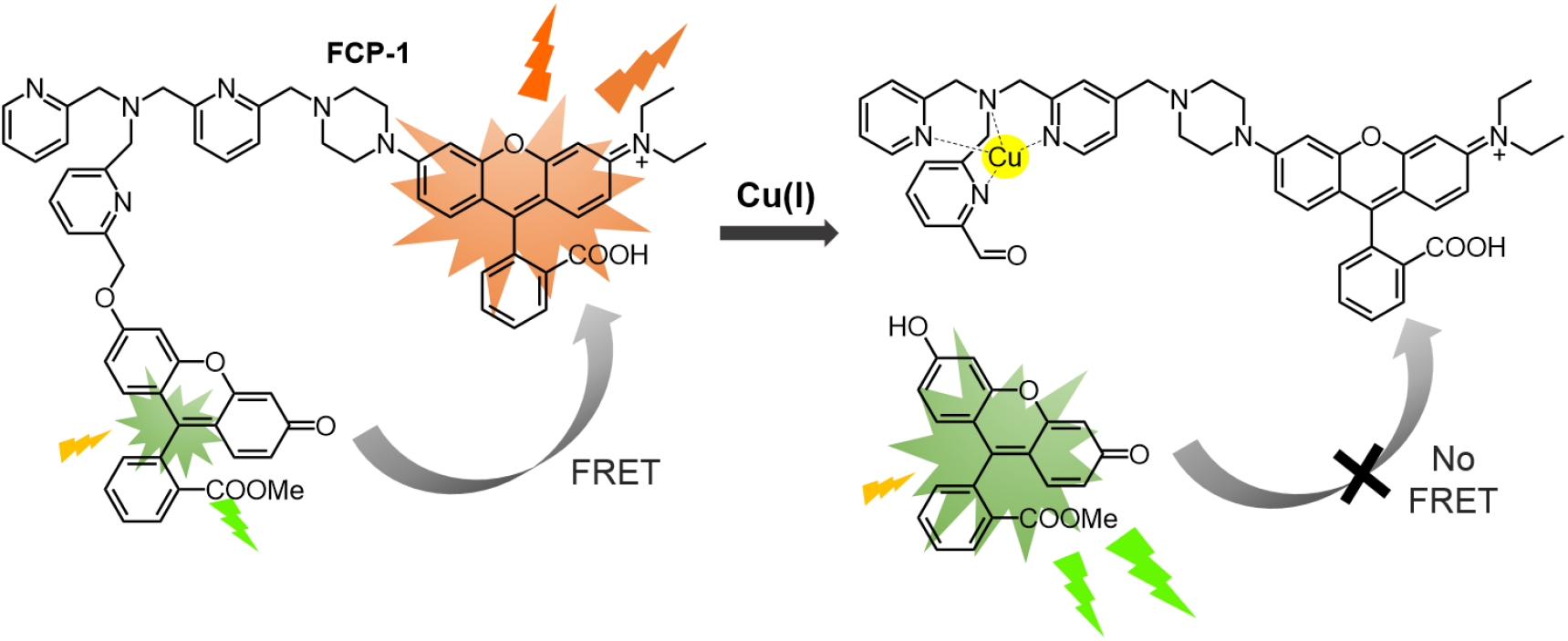
Chemical structure and design of FRET Copper Probe-1, **FCP-1**

**Scheme 2.**
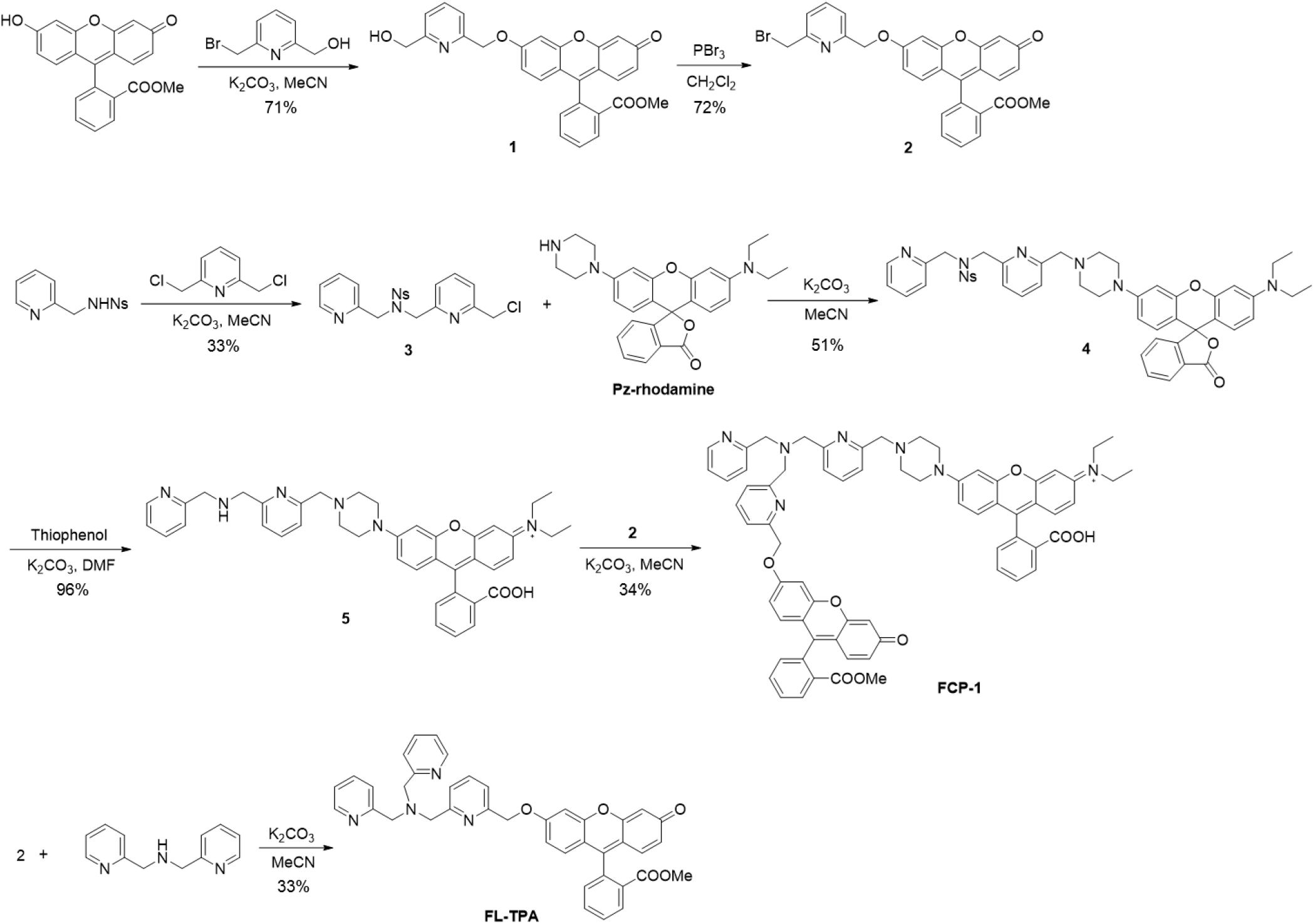
Synthesis of **FCP-1** and control probe **FL-TPA**

