## Supplementary information for "Activity-Based Ratiometric FRET Probe Reveals Oncogene-Driven Changes in Labile Copper Pools Induced by Altered Glutathione Metabolism"

**This PDF file includes:**

Supplementary text
Figs. S1 to S22
References for SI reference citations

### Supplementary Information Text

**Materials and Reagents.** Fluorescein, phosphorous tribromide, 2-(aminomethyl)pyridine and thiophenol were purchased from Sigma-Aldrich. 6-(Bromomethyl)-2-pyridinemethanol and 2,6-bis(chloromethyl)pyridine were purchased from TCI America. 2-(4-Diethylamino-2-hydroxybenzoyl)benzoic acid was purchased from AK Scientific. 3-(1-Piperazinyl)phenol was purchased from Alfa Aesar. For the materials and reagents for *in vitro* and biological assays, copper(II) chloride and bathocuproine disulfonate (BCS) were purchased from Sigma-Aldrich. Dulbecco's modified Eagle's medium (DMEM; high glucose and no phenol red), GlutaMAX, fetal bovine serum (FBS) and phosphate-buffered saline (PBS) were purchased from Thermo Fisher Scientific. Fluorescein methyl ester (1), 4-nitro-*N*-(2-pyridinylmethyl)-benzenesulfonamide (2) and *N*-[9-(2-carboxyphenyl)-6-(1-piperazinyl)-3*H*-xanthen-3-ylidene]-*N*-ethyl-ethanaminium (3) were synthesized according to literature methods. All other reagents were of analytical grade and were used without further purification. Milli-Q water was used in all experiments unless otherwise stated.

**Physical Measurements and Instrumentation.** $^{1}$H NMR and $^{13}$C{$^{1}$H} spectra were collected at 25 °C on Bruker AVB-400, AVQ-400 and AV-300 at the College of Chemistry NMR Facility at the University of California, Berkeley. All chemical shifts are reported in the standard δ notation of parts per million relative to residual solvent peak as an internal reference. Splitting patterns are indicated as follows: br, broad; s, singlet; d, doublet; t, triplet; m, multiplet; dd, doublet of doublets. Low-resolution electrospray mass spectra were recorded on a LC-MS (Agilent Technology 6130, Quadrupole LC/MS). High-resolution mass spectra were collected at the College of Chemistry Mass Spectrometry Facility at the University of California, Berkeley. FCP-1 was characterized by LC-MS (Agilent Technology 6130, Quadrupole LC/MS) coupled with photodiode array for detection (λ = 488 nm). UV-vis absorption spectra were recorded using a Varian Cary 50 spectrophotometer, and fluorescence spectra were recorded using a Photon Technology International Quanta Master 4 L-format scan spectrofluorometer equipped with an LPS-220B 75-W Xenon lamp and power supply, A-1010B lamp housing with integrated igniter, switchable 814 photocounting/analog photomultiplier detection unit, and MD5020 motor driver. Confocal microscopy images were recorded on a Zeiss laser scanning microscope 710 with a 20× or 63× oil-immersion objective lens using Zen 2009 software (Carl Zeiss). ICP-MS were recorded on a Thermo Fisher iCAP-Qc ICP-MS in Kinetic Energy Discrimination (KED) mode with the He flow set to 4.426 mL/min. Measurements were normalized to a standard curve of known copper concentrations doped with 20 ppb Ga. The standard curve was diluted from CMS-5 (Inorganic Ventures).

### Synthesis

**2-[3-(2-Hydroxymethyl-6-pyridylmethoxy)-6-hydroxyl-9*H*-xanthen-9-yl]benzoic acid methyl ester (1).** Fluorescein methyl ester (44.2 mg, 0.13 mmol) and 6-(Bromomethyl)-2-pyridinemethanol (28.4 mg, 0.14 mmol) were dissolved in dry acetonitrile (30 mL). Potassium carbonate (97.0 mg, 0.70 mmol) was added to the solution, and the reaction mixture was heated at 85 °C overnight. The solution mixture was cooled to room temperature, and any undissolved solid was filtered off. The filtrate was evaporated under reduced pressure, and the product was purified by column chromatography on silica gel using ethyl acetate-methanol (25:1, v/v) as the eluent, yielding the desired product as an orange-yellow solid (42.5 mg, 71%). $^{1}$H NMR ($CDCl_3$, 300 MHz): $\delta$ 8.20-8.25 (1H, m), 7.62-7.76 (3H, m), 7.37 (1H, d, $J$ = 7.6 Hz), 7.22-7.32 (2H, m), 7.01 (1H, d, $J$ = 2.3 Hz), 6.78-6.92 (3H, m), 6.51 (1H, dd, $J$ = 1.9 Hz), 6.42 (1H, d, $J$ = 1.9 Hz), 5.28 (2H, s), 4.79 (2H, s), 3.63 (3H, s). $^{13}$C{$^{1}$H} NMR ($CDCl_3$, 75 MHz) $\delta$ 185.9, 165.7, 162.8, 159.3, 159.0, 154.8, 154.2, 150.2, 137.8, 134.7, 132.9, 131.3, 130.7, 130.4, 130.1, 129.8, 129.1, 120.0,

119.99, 117.9, 115.4, 113.6, 105.9, 101.8, 71.2, 64.2, 52.5. LRMS (ESI) m/z $[M+K^+]^+$ calcd for $C_{28}H_{21}NO_6K$: 506.1; found: 506.4.

**2-[3-(2-Bromomethyl-6-pyridylmethoxy)-6-hydroxyl-9*H*-xanthen-9-yl]benzoic acid methyl ester (2). 1** (42.5 mg, 0.09 mmol) was dissolved in dry dichloromethane (30 mL), and phosphorous tribromide (5.12 μL, 0.5 mmol) in dry dichloromethane (10 mL) was added dropwise to the solution at 0 °C. The reaction mixture was then allowed to warm up to room temperature and stir for 3 h. The reaction was quenched by sat. $NaHCO_3$(aq) solution, and the aqueous layer was extracted with dichloromethane three times. The volatile organic solvent was evaporated under reduced pressure, and the product was purified by column chromatography on silica gel using ethyl acetate as the eluent, yielding the desired product as an orange-yellow solid (34.8 mg, 72%). $^1H$ NMR ($CDCl_3$, 300 MHz): $\delta$ 8.22-8.26 (1H, m), 7.63-7.78 (3H, m), 7.38-7.44 (2H, m), 7.27-7.32 (1H, m), 7.04 (1H, d, *J* = 2.3 Hz), 6.79-6.92 (3H, m), 6.52 (1H, dd, *J* = 1.9 Hz), 6.44 (1H, d, *J* = 1.9 Hz), 5.27 (2H, s), 4.56 (2H, s), 3.63 (3H, s). $^{13}C\{^1H\}$ NMR ($CDCl_3$, 75 MHz) $\delta$ 185.9, 165.7, 162.7, 159.0, 156.8, 155.9, 154.2, 150.0, 138.2, 134.7, 132.8, 131.3, 130.7, 130.4, 130.3, 130.2, 129.8, 129.1, 123.0, 120.9, 118.0, 115.4, 113.5, 106.0, 101.9, 71.2, 52.6, 33.6. LRMS (ESI) m/z $[M+H^+]^+$ calcd for $C_{28}H_{21}BrNO_5$: 530.1; found: 530.5.

***N*-[(6-Chloromethyl-2-pyridinyl)methyl]-4-nitro-*N*-(2-pyridinylmethyl)-benzenesulfonamide (3).** 4-nitro-*N*-(2-pyridinylmethyl)-benzenesulfonamide (300 mg, 1.02 mmol) and 2,6-bis(chloromethyl)pyridine (180.1 mg, 1.02 mmol) were dissolved in dry acetonitrile (30 mL). Potassium carbonate (212.1 mg, 1.53 mmol) was added to the solution, and the reaction mixture was heated at 85 °C overnight. The solution mixture was cooled to room temperature, and any undissolved solid was filtered off. The filtrate was evaporated under reduced pressure, and the product was purified by column chromatography on silica gel using hexane-ethyl acetate (1:1, v/v) as eluent, yielding the desired product as a pale yellow oil (146.9 mg, 33%). $^1H$ NMR ($CDCl_3$, 300 MHz): $\delta$ 8.32-8.37 (1H, m), 8.19-8.26 (2H, m), 7.92-7.99 (2H, m), 7.35 (1H, d, *J* = 7.8 Hz), 7.26 (1H, d, *J* = 7.6 Hz), 7.22 (1H, d, *J* = 7.7 Hz), 7.09-7.15 (1H, m), 4.66 (2H, s), 4.63 (2H, s), 4.37 (2H, s). $^{13}C\{^1H\}$ NMR ($CDCl_3$, 75 MHz) $\delta$ 156.3, 155.6, 155.3, 149.8, 149.3, 146.0, 137.8, 136.8, 128.7, 124.0, 122.8, 122.1, 121.9, 53.3, 52.7, 46.4. LRMS (ESI) m/z $[M+H^+]^+$ calcd for $C_{19}H_{18}ClN_4O_4S$: 433.1; found: 433.5.

**Compound 4**. *N*-[9-(2-carboxyphenyl)-6-(1-piperazinyl)-3*H*-xanthen-3-ylidene]-*N*-ethyl-ethanaminium (188.7 mg, 0.41 mmol) and **3** (196.8 mg, 0.45 mmol) were dissolved in dry acetonitrile (30 mL). Potassium carbonate (212.1 mg, 1.53 mmol) was added to the solution, and the reaction mixture was heated at 85 °C overnight. The solution mixture was cooled to room temperature, and any undissolved solid was filtered off. The filtrate was evaporated under reduced pressure, and the product was purified by column chromatography on alumina using ethyl acetate-methanol (9:1, v/v) as eluent, yielding the desired product as pale pink film (180.1 mg, 51%). $^1H$ NMR ($CDCl_3$, 300 MHz): $\delta$ 8.34 (1H, d, *J* = 4.7 Hz), 8.24 (2H, d, *J* = 8.9 Hz), 7.94-8.03 (3H, m), 7.53-7.67 (4H, m), 7.34 (1H, d, *J* = 7.9 Hz), 7.30 (1H, d, *J* = 7.7 Hz), 7.09-7.25 (3H, m), 6.67 (1H, d, *J* = 2.0 Hz), 6.51-6.64 (3H, m), 6.42 (1H, d, *J* = 2.4 Hz), 6.29-6.37 (1H, m), 4.67 (2H, s), 4.66 (2H, s), 3.46 (2H, s), 3.34 (4H, q, *J* = 7.1 Hz), 3.18-3.28 (4H, br), 2.51-2.58 (4H, br), 1.15 (6H, t, *J* = 7.0 Hz). $^{13}C\{^1H\}$ NMR ($CDCl_3$, 75 MHz) $\delta$ 169.9, 158.3, 155.7, 155.0, 153.3, 153.2, 152.9, 152.8, 149.8, 149.6, 149.3, 146.1, 137.3, 136.8, 134.7, 129.4, 129.0, 128.8, 127.5, 124.8, 124.1, 124.0, 122.8, 122.7, 122.0, 121.0, 111.5, 109.8, 108.2, 105.5, 102.0, 97.6, 84.9, 64.2, 60.5, 53.3, 53.2, 53.1, 48.2, 44.5, 12.6. LRMS (ESI) m/z $[M+H^+]^+$ calcd for $C_{47}H_{46}N_7O_7S$: 852.3; found: 852.6.

**Compound 5. 4** (42.6 mg, 0.05 mmol) was dissolved in dry dimethylformamide (2 mL). Potassium carbonate (31.7 mg, 0.23 mmol) was added to the solution, followed by the addition of thiophenol (51.3 μL, 0.43 mmol). The solution mixture was stirred at room temperature overnight. The solvent was evaporated under vacuum, and the crude product was re-dissolved in ethyl acetate. Any undissolved solid was filtered off, and the organic solvent was evaporated under reduced pressure. The crude product was dissolved in minimum amount of dichloromethane with 1 vol% trifluoroacetic acid, and precipitated by adding large volume of diethyl ether. The solid was further washed by diethyl ether twice, and dried under vacuum to yield the desired product as purple film (37.5 mg, 96%). $^{1}$H NMR ($CDCl_3$, 400 MHz): $\delta$ 8.62 (1H, d, $J$ = 4.8 Hz), 8.37 (1H, d, $J$ = 7.8 Hz), 8.00 (1H, t, $J$ = 7.8 Hz), 7.79-7.91 (3H, m), 7.37-7.60 (7H, m), 7.13-7.30 (4H, m), 7.04 (1H, d, $J$ = 1.6 Hz), 4.67 (2H, s), 4.60 (2H, s), 4.57 (2H, s), 4.01-4.11 (4H, br), 3.73 (4H, q, $J$ = 7.1 Hz), 3.61-3.68 (4H, br), 1.33 (6H, t, $J$ = 7.0 Hz). $^{13}$C{$^{1}$H} NMR ($CDCl_3$, 100 MHz) $\delta$ 168.0, 160.1, 158.7, 158.3, 157.3, 153.3, 152.5, 151.0, 150.6, 140.5, 138.9, 135.1, 134.0, 133.0, 132.6, 132.2, 131.6, 131.4, 130.2, 125.4, 125.1, 124.4, 124.3, 117.0, 116.4, 115.9, 115.5, 100.1, 97.3, 66.9, 60.8, 52.8, 51.7, 51.4, 47.2, 45.0, 12.8. LRMS (ESI) m/z $[M]^+$ calcd for $C_{41}H_{43}N_6O_3$: 667.3; found: 667.9.

**FCP-1.** Compounds **2** (11.0 mg, 0.021 mmol) and **5** (20.2 mg, 0.021 mmol) were dissolved in dry acetonitrile (30 mL). Potassium carbonate (20.1 mg, 0.145 mmol) was added to the solution, and the reaction mixture was heated at 85 $^{o}$C overnight. The solution mixture was cooled to room temperature, and any undissolved solid was filtered off. The filtrate was evaporated under reduced pressure, and the product was purified by prep-HPLC. The mobile phase was a mixture of 0.1% formic acid in water (phase A) and 0.1% formic acid in methanol (phase B). Separation was achieved by using: 50% phase B at $t$ = 0 min; 50–100% phase B from 0 to 40 min; 100% phase B from 40 to 100 min. The flow rate was 1.5 mL/min and the injection volume was 2 mL. The solvent mixture was evaporated under vacuum, yielding the desired product as red film (7.8 mg, 34%). $^{1}$H NMR ($CDCl_3$, 300 MHz): $\delta$ 8.54 (1H, d, $J$ = 5.4 Hz), 8.23 (1H, d, $J$ = 6.9 Hz), 7.99 (1H, d, $J$ = 7.7 Hz), 7.55-7.75 (10H, m), 7.52 (1H, s), 7.29-7.36 (3H, m), 7.12-7.20 (2H, m), 7.04 (1H, d, $J$ = 2.3 Hz), 6.78-6.90 (3H, m), 6.66 (1H, m), 6.50-6.61 (4H, m), 6.42 (2H, d, $J$ = 2.1 Hz), 6.31-6.35 (1H, m), 5.26 (2H, s), 3.92 (2H, s), 3.91 (4H, s), 3.71 (2H, s), 3.63 (3H, s), 3.35 (4H, q, $J$ = 7.0 Hz), 3.22-3.27 (4H, br), 2.63-2.68 (4H, br), 1.18 (6H, t, $J$ = 7.0 Hz). $^{13}$C{$^{1}$H} NMR ($CDCl_3$, 100 MHz) $\delta$ 185.9, 169.9, 165.7, 163.0, 159.5, 159.2, 158.9, 157.0, 155.1, 154.3, 153.3, 153.1, 152.9, 152.8, 149.8, 149.1, 137.6, 137.2, 136.8, 134.6, 132.9, 131.3, 130.7, 130.5, 130.0, 129.9, 129.4, 129.2, 129.1, 128.9, 127.8, 125.0, 124.3, 123.2, 122.5, 122.4, 122.0, 121.7, 119.9, 117.8, 115.4, 113.8, 111.7, 110.0, 108.4, 105.9, 105.8, 102.0, 101.9, 97.7, 71.5, 64.1, 60.3, 60.1, 52.9, 52.6, 48.0, 44.6, 12.7. HRMS (ESI) m/z $[M]^+$ calcd for $C_{69}H_{62}N_7O_8$: 1116.4660; found: 1116.4660.

**FL-TPA.** Compound **2** (40 mg, 0.075 mmol) and di-(2-picolyl)amine (15.0 mg, 0.075 mmol) were dissolved in dry acetonitrile (30 mL). Potassium carbonate (57.3 mg, 0.41 mmol) was added to the solution, and the reaction mixture was heated at 85 $^{o}$C overnight. The solution mixture was cooled to room temperature, and any undissolved solid was filtered off. The filtrate was evaporated under reduced pressure, and the product was purified by column chromatography on alumina using ethyl acetate-methanol (49:1, v/v) as eluent, yielding the desired product as yellow-orange film (15.0 mg, 31%).$^{1}$H NMR ($CDCl_3$, 300 MHz): $\delta$ 8.54 (2H, d, $J$ = 6.0 Hz), 8.21-8.26 (1H, m), 7.53-7.74 (8H, m), 7.28-7.36 (2H, m), 7.15 (2H, t, $J$ = 5.7 Hz), 7.04 (1H, d, $J$ = 2.5 Hz), 6.79-6.91 (3H, m), 6.53 (1H, dd, $J$ = 2.0 and 4.5 Hz), 6.42 (1H, d, $J$ = 1.8 Hz), 5.26 (2H, s), 3.92 (2H, s), 3.90 (4H, s), 3.63 (3H, s). $^{13}$C{$^{1}$H} NMR ($CDCl_3$, 100 MHz) $\delta$ 185.7, 165.7, 162.9, 159.0, 156.7, 155.4, 154.3, 150.2, 148.8, 137.9, 137.5, 134.7, 132.8, 131.3, 130.7, 130.4, 130.3, 130.1, 129.8, 129.1, 124.0, 123.0, 120.5, 117.9, 115.4, 113.7, 105.9, 101.9, 71.3, 59.4, 59.2, 53.6, 52.5. HRMS (ESI) m/z $[M+H]^+$ calcd for $C_{40}H_{33}N_4O_5$: 649.2451; found: 649.2454.

**LC-MS characterization of FCP-1.** FCP-1 were dissolved in a $H_2O$/MeOH solution mixture, and an aliquot was taken for LC-MS analysis. Separation was achieved by gradient elution from 5-100% MeOH in water (constant 0.1 vol% formic acid) over 8 min, isocratic with 100% MeOH from 8 to 12 min and returned to initial conditions and equilibrated for 3 minutes. The LC chromatograms were recorded by monitoring absorption at 488 nm.

**UV-vis absorption and emission measurements.** FCP-1 stock solution in DMSO (5 mM) was prepared, aliquoted and stored at −20 °C. For each spectroscopic measurement, a fresh aliquot was diluted to 500 μM by DMSO and further diluted with PBS containing 2 mM GSH/PEG-400 mixture (3:2, v/v) to prepare the sensing solution with 5 μM of FCP-1**.** The sensing solution was pre-incubated in a water bath at 37 °C, and then $[Cu(CH_3CN)](PF_6)$ in MeCN or other metal ions in $H_2O$ was added. The solution mixture was kept at 37 °C and taken out for UV-vis absorption or emission spectroscopic measurements in quartz cuvette at predetermined time intervals. Emission spectra were corrected using coumarin 153 as secondary emission standard (4). Photoluminescence quantum yield (PLQY) was determined using fluorescein in 0.1 M NaOH solution (0.95; $\lambda_{ex}$ = 496 nm). FRET efficiency from the fluorescein moiety to rhodamine moiety of FCP-1 was estimated by [1 – (PLQY of FCP-1/PLQY of FL-TPA)].

**Inductively coupled plasma-mass spectrometry (ICP-MS)**. HeLa cells were plated on 6-well plates. At *ca.* 40% confluency, the cells were treated with solvent control, different concentrations of $CuCl_2$ or BCS (100 μM) in complete medium for 24 h. The cells were then washed with 20 mM HEPES buffer (pH 7.4) three times and then digested in concentrated nitric acid (100 mg/mL $HNO_3$, BDH Aristar Ultra) overnight and heated up at 90 ºC for 2 h in 1.5 mL tubes (Sarstedt) with small holes poked in the caps. Once cooled to room temperature, samples were diluted into 2% $HNO_3$ and doped with a gallium internal standard (Inorganic Ventures, 20 ppb final concentration). The copper and phosphorous content was determined by measuring $^{63}$Cu and $^{31}$P using a Thermo Fisher iCAP-Qc ICP-MS in Kinetic Energy Discrimination (KED) mode with the He flow set to 4.426 mL/min. Measurements were normalized to a standard curve of known copper concentrations doped with 20 ppb Ga. The standard curve was diluted from CMS-5 (Inorganic Ventures).

Similarly, mouse embryonic fibroblasts (MEF) *Ctr1*$^{+/+}$ and *Ctr1*$^{-/-}$ cells were grown on 6-well plates. At *ca*. 70% confluency, the cells were washed with PBS three times, digested in concentrated nitric acid overnight, heated up at 90 ºC for 2 h and then diluted into 2% $HNO_3$ and doped with a gallium internal standard. The solutions were then assayed for copper and phosphorous content by measuring $^{63}$Cu and $^{31}$P using a Thermo Fisher iCAP-Qc ICP-MS.

**Cell culture.** Following cell cultures were grown in the UC Berkeley Tissue Culture Facility: Human embryonic kidney (HEK 293T) cells were maintained in Dulbecco’s Modified Eagle Medium (DMEM) supplemented with 10 vol% fetal bovine serum (FBS), 1 vol% GlutaMax (Gibco) and 1 vol% non-essential amino acids (NEAA, Gibco). *Ctr1*$^{+/+}$ and *Ctr1*$^{-/-}$ MEF cells were maintained in DMEM with 10 vol% FBS, 1 vol% non-essential amino acids, 1 vol% sodium pyruvate and 0.1 vol% 2-mercaptoethanol or DMEM supplemented with 10vol% FBS and 1x Penicillin-Streptomycin. Human cervical epithelial carcinoma (HeLa) was maintained in DMEM medium supplemented with 1 vol% GlutaMAX and 10 vol% FBS. All cells were incubated in 5% $CO_2$ humidified air and subcultured at 80% confluence.

For MEFs (immortalized with a plasmid encoding the SV40-T-Ag gene), they were maintained in Dulbecco’s Modified Eagle Medium (DMEM) supplemented with 10 vol% fetal bovine serum (FBS), 1x Penicillin/ Streptomycin (PS) as described previously. Immortalized MEFs were stably infected with retroviruses derived from pBABE or pWZL (*see* plasmids below) or lentiviruses derived from pLKO.1 (*see* plasmids below) using established protocols. MEFs stably expressing non-targeting shRNA (Scr), *Gclc* shRNA, *Gsr* shRNA, *BRAF*$^{V600E}$ cDNA, *KRAS*$^{G12D}$ cDNA, or *BRAF*$^{V600E}$ cDNA with *Mek1* shRNA and *MEK1* cDNA, or

*BRAF*$^{V600E}$ cDNA with *Mek1* shRNA and *MEK1*$^{CBM}$ cDNA were maintained in DMEM supplemented with 10 vol% FBS), 1x PS, and 2 µg/mL puromycin and/or 5 µg/mL blasticidin.

**Plasmids.** pBABEpuro-HA-KRAS$^{G12D}$ (Addgene plasmid #58902), pBABEpuro-MYC-HIS-BRAF$^{V600E}$ (Addgene plasmid #15269), pWZLblasti-HA-MEK1 (Addgene plasmid #53161), and pWZLbalsti-HA-MEK1$^{CBM}$ (5) were used. The TRC cloning vector pLKO.1 expressing the mouse *Gclc* shRNA target sequence 5'-CCTCTCATACAAACTAGACTT-3', the mouse *Gsr* shRNA target sequence 5'-CCCAAATTCTAAGGGCCTGAA-3', or the mouse Mek1 shRNA target sequence 5'-CCTGGAGATCAAACCCGCAAT -3' was obtained from the High-Throughput Screening Core at the University of Pennsylvania.

**RNA extraction and Reverse transcriptase quantitative PCR.** For RT-qPCR, RNA was purified from MEFs and reverse transcribed to cDNA using standard protocols and then quantified utilizing Taqman probes: Mm00558247_m1 to detect mouse *Ctr1*, Mm00437663_m1 to detect mouse *Atp7a*, Mm00439154_m1 to detect mouse *Gsr*, Mm00802655_m1 to detect mouse *Gclc*, Mm_01277042_m1 to detect mouse TATA binding protein (*Tbp*), Mm01318743_m1 to detect mouse *Hprt* and Mm02619580_g1 to detect mouse beta Actin (*Actb*) using the ViiA 7 Real-Time PCR System. Relative mRNA expression levels were normalized to *Tbp*, *Actb*, *Hprt* and analyzed using comparative delta-delta CT method and represented as a fold change.

**Immunoblot analysis.** Indicated cell lines were washed with cold PBS and lysed with cold RIPA buffer containing 1X EDTA-free Halt™ protease and phosphatase inhibitor cocktail halt protease and phosphatase inhibitors (Thermo Scientific). The protein concentration was determined by BCA Protein Assay (Pierce) using BSA as a standard. Equal amount of lysates were resolved by SDS-PAGE using standard techniques, and protein was detected with the following primary antibodies: mouse anti-γ-GCSc (1:1000, sc-390811, Santa Cruz Biotechnology), rabbit anti-glutathione reductase (1:1000, ab16801, Abcam), mouse anti- β-Actin (1:10000, 3700S, Cell Signaling), or rabbit anti-CCS (1:1000, sc-20141, Santa Cruz Biotechnology) followed by detection with one of the horseradish peroxidase conjugated secondary antibodies: goat anti-rabbit IgG (1:5000, 7074, Cell Signaling) or goat anti-mouse IgG (1:5000, 7076, Cell Signaling), using SignalFire (Cell Signaling) or SignalFire Elite ECL (Cell Signaling) detection reagents. The fold change in the total protein to actin was measured in Image Studio Lite (LI-CORE Biosciences) software by boxing each band per representative image using the rectangular selection tool and calculating the total signal of the band in pixels. The ratio of total protein to actin pixel intensity for each experimental condition was normalized to control and graphed as average fold change.

**Total glutathione levels and reduced/oxidized glutathione ratios.** MEFs were seeded at 20,000 cells/well in 96-well, white walled, flat clear bottom plates. Twenty-four hours after seeding, MEFs were treated with vehicle, 1 mM BSO or 0.1 mM BCNU for 4 hours and then assayed for totally glutathione levels or GSH/GSSG ratios. Total glutathione levels and the GSH/GSSG ratios were measured using the GSH/GSSG-Glo™ Assay kit from Promega (V6611) following the manufacturer's protocol for adherent cells. Total glutathione measurements and GSH/GSSG ratios were measured in four independent experiments assayed in triplicate. Statistical analysis of total glutathione and GSH/GSSG ratio was analyzed using a one-way ANOVA followed by a Dunnett's multi-comparisons test in Prism7 (GraphPad).

**Confocal fluorescence microscopy imaging.** For confocal fluorescence imaging experiments of HEK293T, MEF *Ctr1*$^{+/+}$ and *Ctr1*$^{-/-}$ and HeLa cells, the cells were plated on 8-well Lab Tek borosilicate chambered coverglass slides (Nunc), and allowed to grow to *ca*. 60% confluency before performing the cell imaging experiments. The confocal imaging was performed with a Zeiss laser scanning microscope 710 with 20× or 63× oil-immersion objective lens using Zen 2009 software (Carl Zeiss). FCP-1 was excited with 458 nm with an Ar laser, and the emissions were collected using a META detector between 465 and 541 nm ($F_{green}$), and between 559 and 710 nm ($F_{orange}$). DRAQ-5 was excited with a 633 nm He Ne laser, and its emission was collected using a META detector between 661 and 759 nm. Image analysis was performed using ImageJ, with ratiometric images generated by the Ratio Plus plugin of ImageJ. For quantification, a threshold value was set as background and the average intensity of the whole image was measured by ImageJ. All experiments were performed in triplicate, and statistical analyses were performed with a two-tailed Student's t-test (MS excel).

For imaging experiments with MEFs stably expressing with *Gclc* shRNA/ *Gsr* shRNA, $BRAF^{V600E}$, $KRAS^{G12D}$, $BRAF^{V600E}$ with Mek1 shRNA and MEK1, or $BRAF^{V600E}$ with Mek1 shRNA and $MEK1^{CBM}$, cells were seeded at 150,000 cells/plate in 35 mm glass insert imaging plates (MatTek) and 48 hours later were imaged. Cells were washed with PBS and media was replaced with Live Cell Imaging Solution (Thermo Scientific) containing 5 μM of FCP-1, 1 μM of CF4, or Ctrl-CF4 for 20 min and imaged on a Zeiss LSM880 laser scanning confocal microscopy system with a 20x dry objective or 63x oil immersion objective lens. Hoechst 33528 was excited with a 405 nm diode laser, and emission was collected between 465 nm and 521 nm. FCP-1 was excited by a 458 nm Ar laser, and emissions were collected between 465 nm and 541 nm ($F_{green}$) and between 559 nm and 710 nm ($F_{orange}$). CF4 and Ctrl-CF4 were excited by a 458 nm Ar laser and emission was collected at 521 nm. FCP-1 image analysis was performed using ImageJ with ratiometric images generated by the Ratio Plus plugin of ImageJ. For quantification, a threshold value was set as background and the average intensity of individual cells was measured by ImageJ. Fluorescence of FCP-1, CF4, or Ctrl-CF4 from single cells was analyzed from at least three independent experiments in which 10 cells per field of view from three independent images. ImageJ was utilized to set thresholds for background, the freehand selection tool was used to select an individual cell, and the measure tool was used to measure the mean intensity, area, and standard deviation. Statistical analysis of normalized FCP-1, CF4, or Ctrl-CF4 fluorescence per cell was performed using a one-way ANOVA followed by a Dunnett's multi-comparisons test in Prism7 (GraphPad).

**Flow Cytometry.** Cells were plated at a density of 150,000 cells/well on 6-well plates (Genesee Scientific, Cat# 25-105) and were treated and collected for flow cytometry 24 hours later. The CellROX Green Flow Cytometry Kit (Invitrogen, Cat# C10492) was used to assess levels of reactive oxygen species in live cells. All cell lines were then treated with 500nM CellROX green reagent for 1 hour before cells were washed with PBS, trypsinized, and assessed by flow cytometry. Data were acquired using the Attune NxT flow cytometry (Thermo Scientific) and analyzed with FlowJo 8.7 (Tree Sar). Statistical analysis of normalized mean fluorescence intensity was performed using a one-way ANOVA followed by a Dunnett's multi-comparisons test in Prism8 (GraphPad).

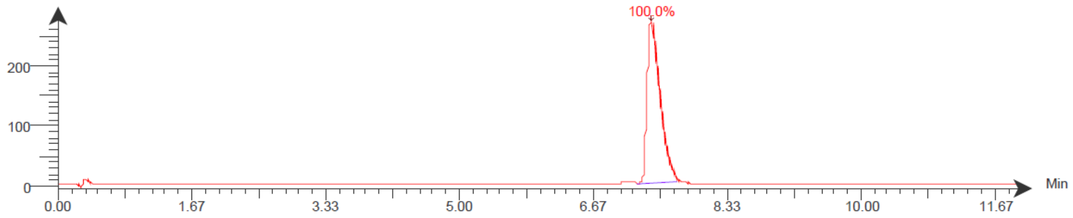


**Fig. S1.** LC trace of FCP-1 monitored at 488 nm.

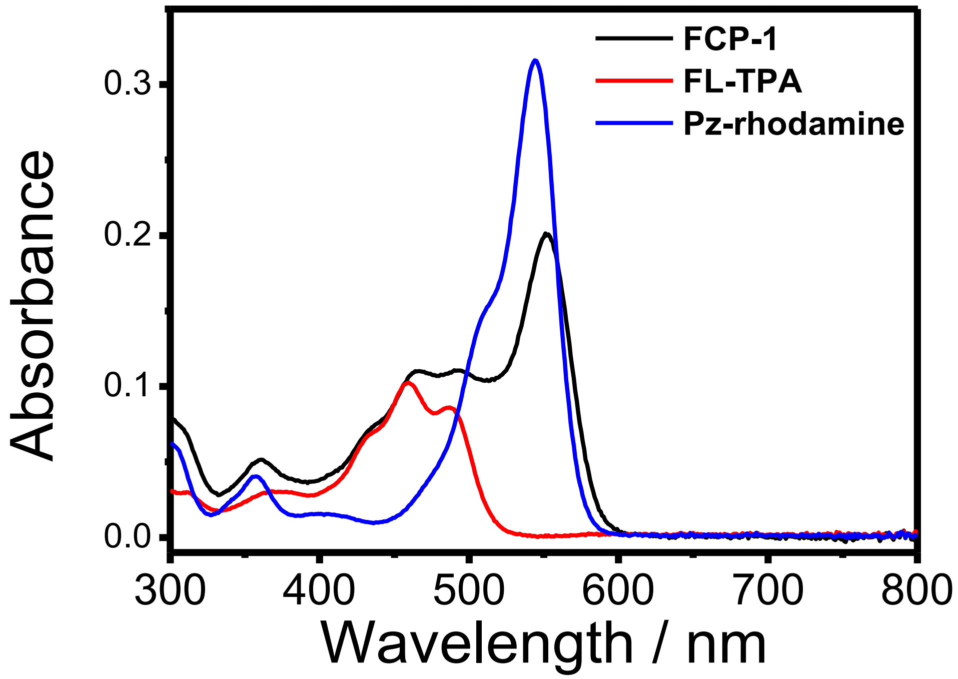


**Fig. S2.** UV-vis absorption spectra of FCP-1, FL-TPA and Pz-rhodamine (5 μM) in PBS containing 2 mM GSH with 40 vol% PEG-400.

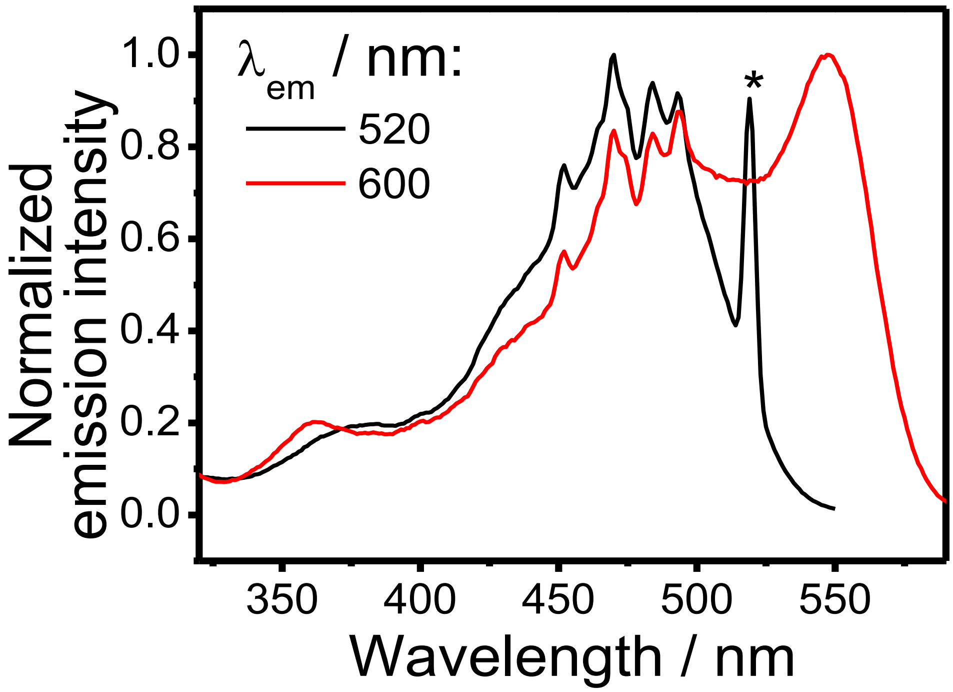


**Fig. S3.** Normalized excitation spectra of FCP-1 (5 μM) in PBS containing 2 mM of GSH with 40 vol% PEG-400. Asterisk denotes scattering from the fluorescence of FCP-1.

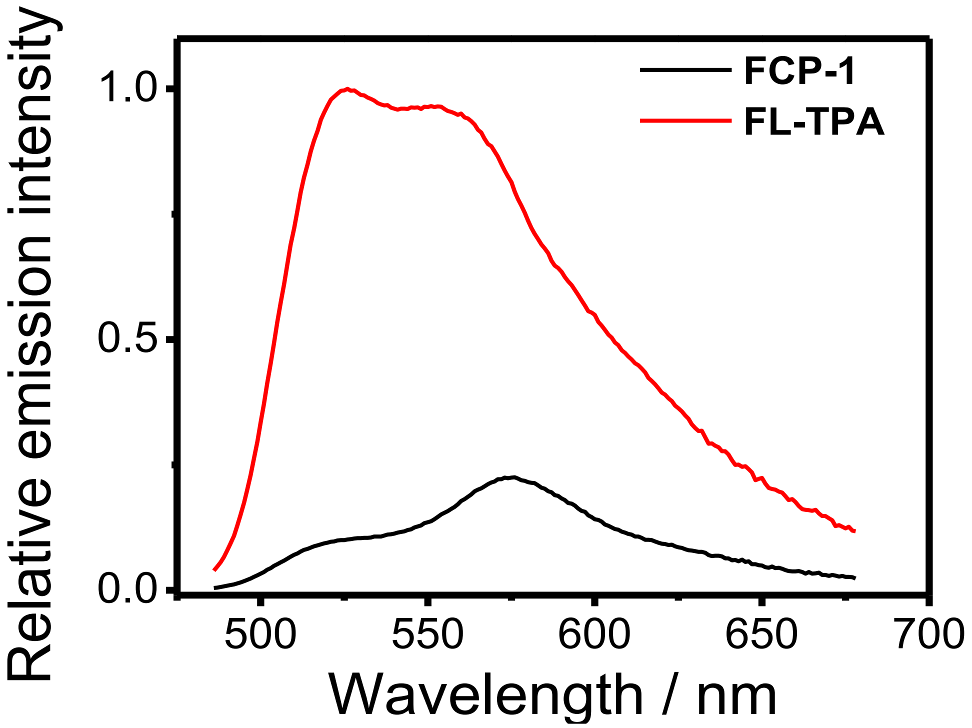


**Fig. S4.** Corrected emission spectra of FCP-1 (5 μM) and FL-TPA (5 μM) in PBS containing 2 mM GSH with 40 vol% PEG-400. $\lambda_{ex}$ = 458 nm.

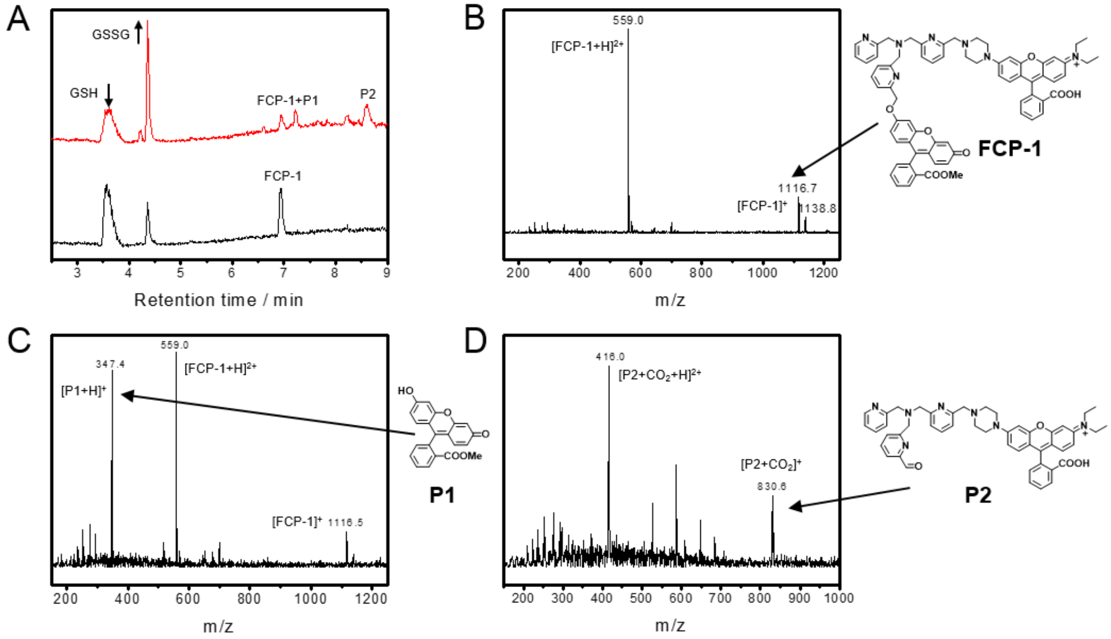


**Fig. S5.** (A) LC-MS chromatograms of PBS solution mixtures (with 2 mM GSH) of FCP-1 (10 µM, black) and FCP-1 (10 µM) after incubation with $[Cu(MeCN)_4](PF_6)$ (100 µM) for 15 min (red). (B) MS spectra at the retention time of 6.93 min in the LC chromatogram of PBS solution mixture of GSH (2 mM) and FCP-1 (10 µM). (C) MS spectra at the retention time of 6.85-7.00 min in the LC chromatogram of PBS solution mixture of GSH (2 mM), FCP-1 (10 µM) and $[Cu(MeCN)_4](PF_6)$ (100 µM).(D) MS spectra at the retention time of 8.61 min in the LC chromatogram of PBS solution mixture of GSH (2 mM), FCP-1 (10 µM) and $[Cu(MeCN)_4](PF_6)$ (100 µM).

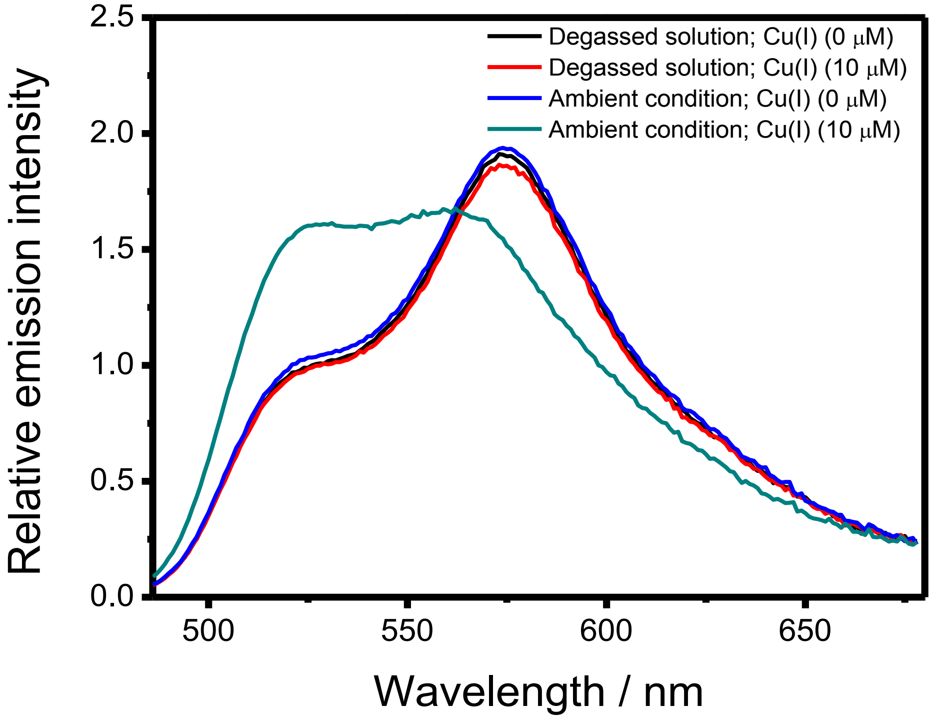


**Fig. S6.** Corrected emission spectra of FCP-1 (5 μM) in PBS containing 2 mM GSH and 40 vol% PEG-400 after incubation with solvent control or Cu(I) (10 μM) for 15 min, respectively, under degassed or ambient condition,. $\lambda_{ex}$ = 458 nm.

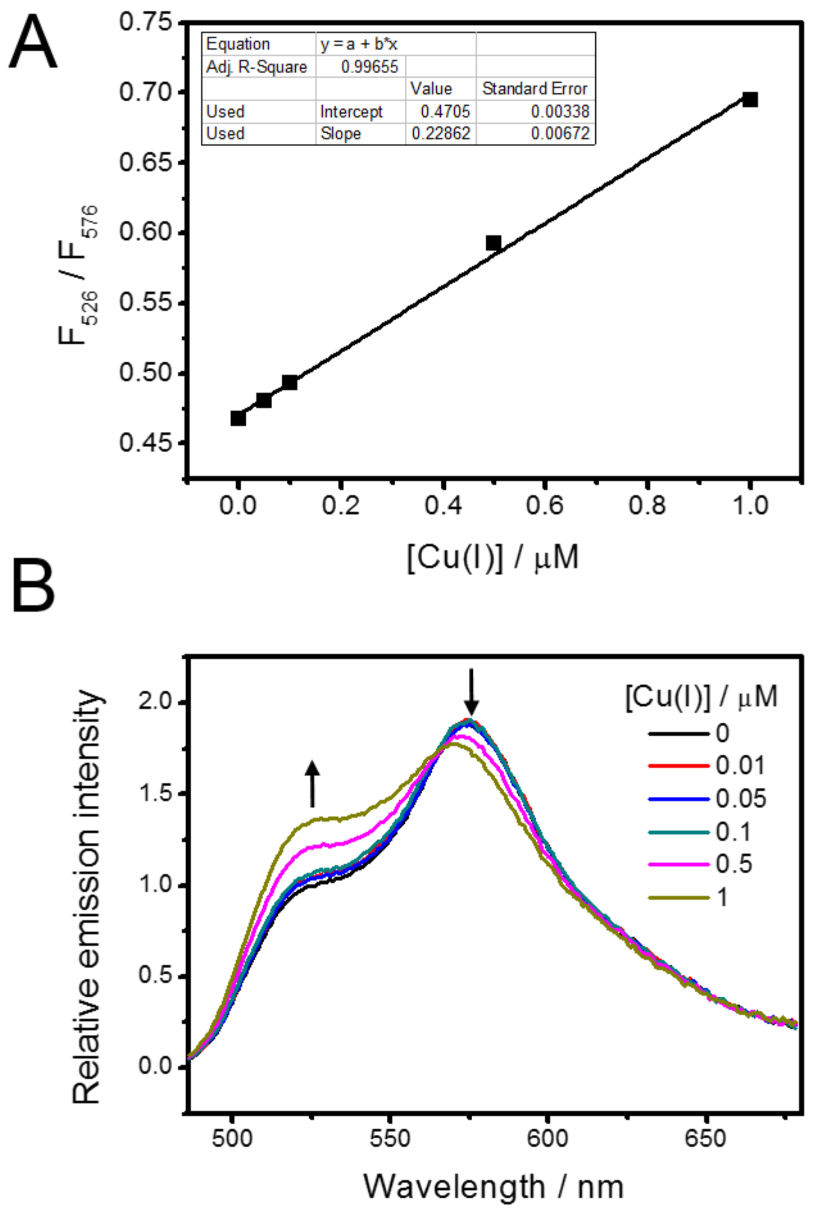


**Fig. S7.** (A) The ratiometric fluorescence change of FCP-1 (5 μM) at 526 and 576 nm ($F_{526}$ / $F_{576}$) and (B) corrected emission spectra of FCP-1 (5 μM), in PBS containing 2 mM GSH and 40 vol% PEG-400 with different concentrations of Cu(I). $\lambda_{ex}$ = 458 nm.

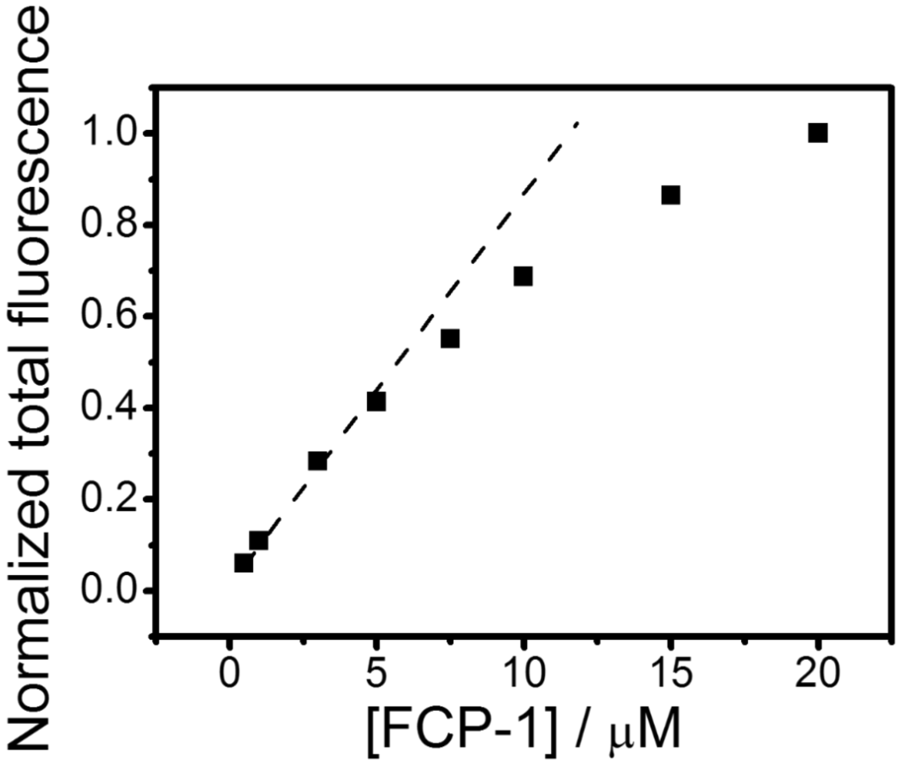


**Fig. S8.** Changes in total fluorescence intensity of FCP-1 in aqueous buffer solution (PBS with 2 mM GSH and 40 vol% PEG-400) at different concentrations. $\lambda_{ex}$ = 458 nm.

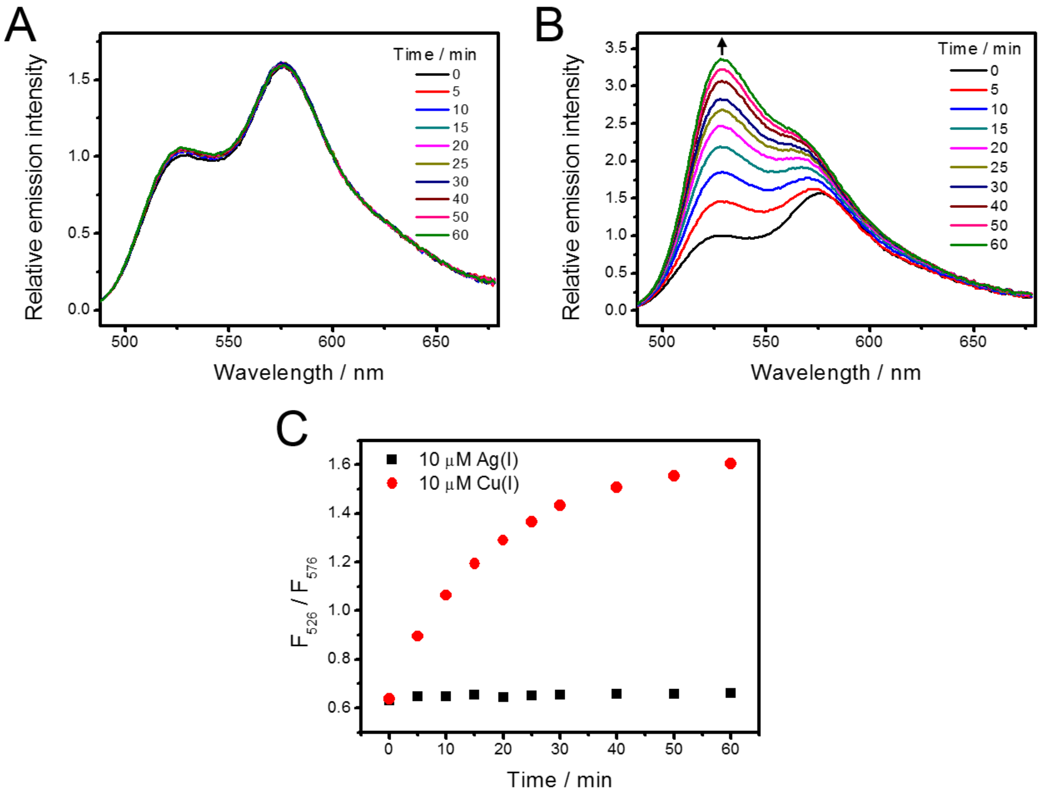


**Fig. S9.** Changes in corrected emission spectra of FCP-1 (5 μM) in $NaH_2PO_4$ buffer solution (50 mM, pH 7.4) containing 2 mM GSH and 40 vol% PEG-400 with (A) Ag(I) (10 μM) and (B) Cu(I) (10 μM) over time. $\lambda_{ex}$ = 458 nm. (C) $F_{526}$ / $F_{576}$ of FCP-1 solution with Ag(I) and Cu(I) respectively

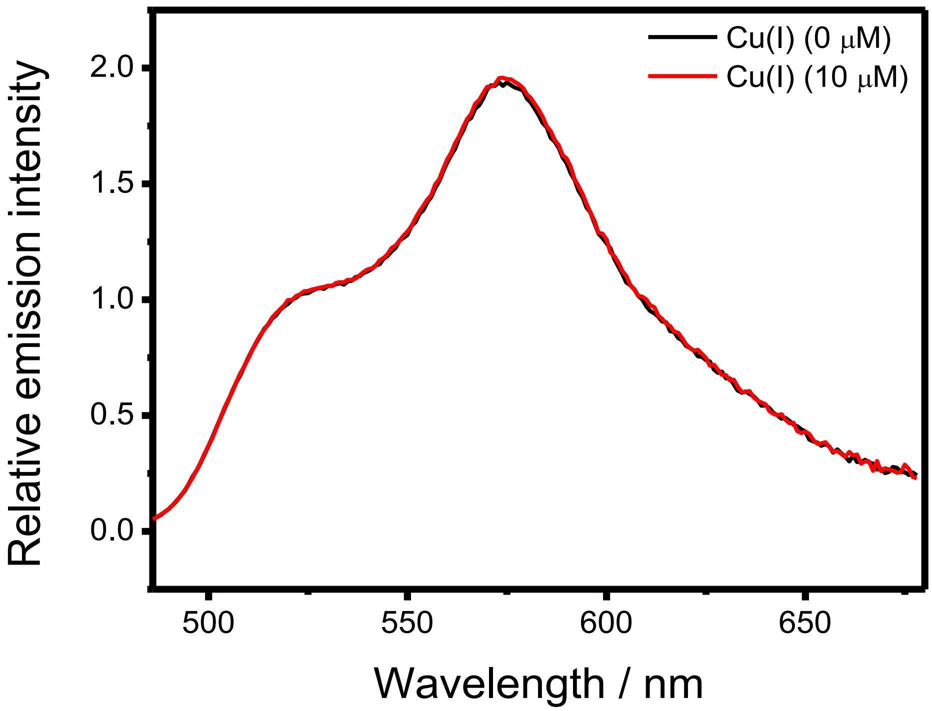


**Fig. S10.** Corrected emission spectra of FCP-1 (5 μM) in PBS containing 40 vol% PEG-400 after incubation with solvent control or Cu(I) (10 μM) for 15 min in the absence of GSH. $\lambda_{ex}$ = 458 nm.

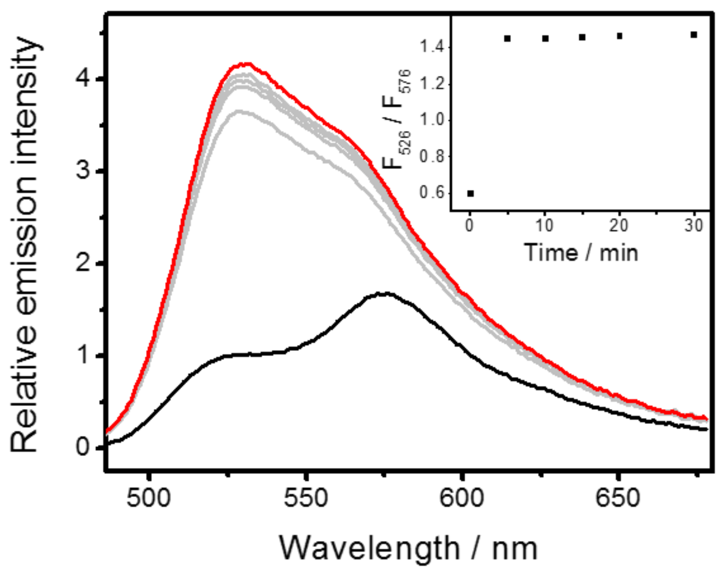


**Fig. S11.** Changes in corrected emission spectra of FCP-1 (5 µM) in PBS containing 2 mM ascorbate and 40 vol% PEG-400 with Cu(I) (10 µM) over time. Inset shows changes in ratiometric emission of FCP-1, $F_{526}$ / $F_{576}$, with Cu(I) over time. $\lambda_{ex}$ = 458 nm.

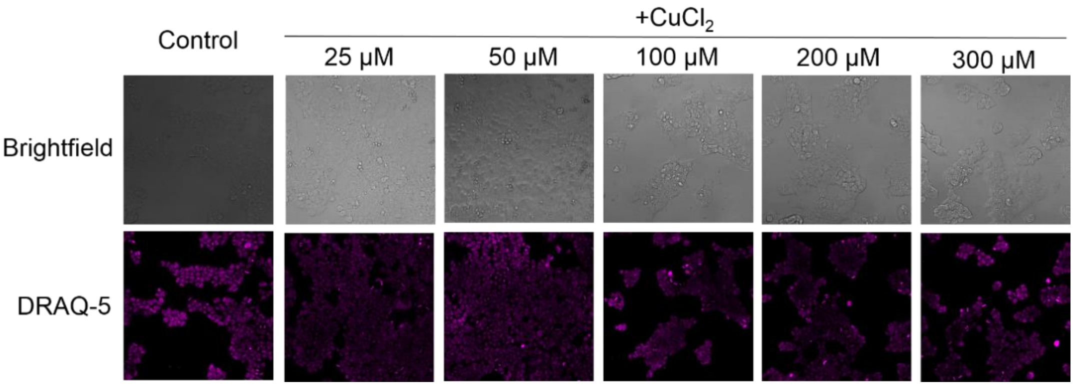


**Fig. S12.** Confocal microscopy images of HEK 293T cells pretreated with solvent vehicle or different concentrations of $CuCl_2$ in complete medium for 18 h, washed with PBS and stained with FCP-1 (5 μM) in DPBS for 45 min. At the last 5-min incubation, a nucleus stain, DRAQ-5 (5 μM), was added to the solution mixture. Without washing, the cells were imaged by confocal fluorescence microscopy.

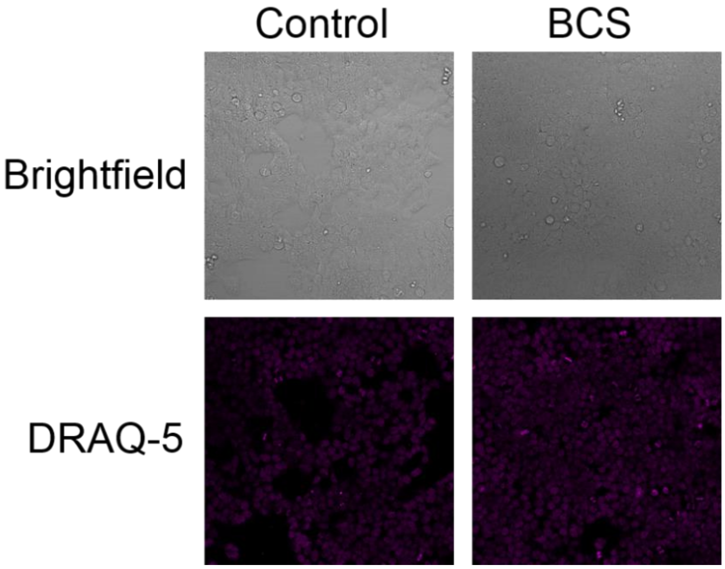


**Fig. S13.** Confocal microscopy images of HEK 293T cells pretreated with solvent vehicle or BCS (100 μM) in complete medium for 18 h, washed with PBS and stained with FCP-1 (5 μM) in DPBS for 45 min. At the last 5-min incubation, a nucleus stain, DRAQ-5 (5 μM), was added to the solution mixture. Without washing, the cells were imaged by confocal fluorescence microscopy.

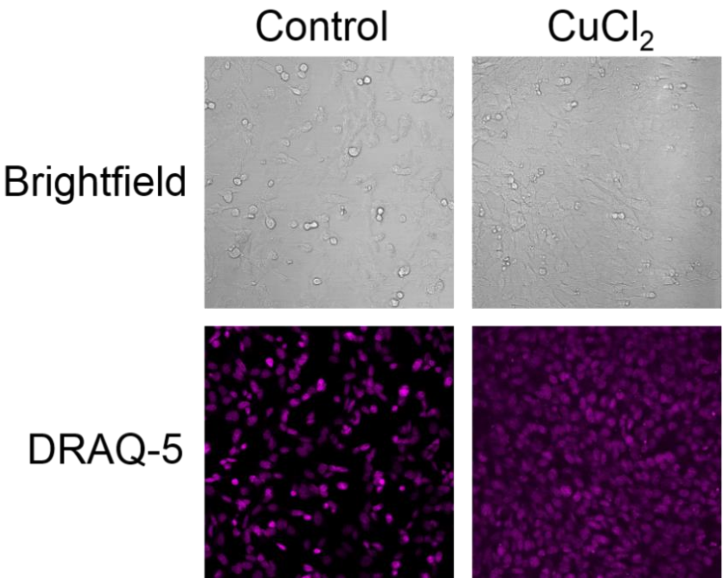


**Fig. S14.** Confocal microscopy images of *Ctr1*$^{+/+}$ MEFs pretreated with solvent vehicle or $CuCl_2$ (300 μM) in complete medium for 8 h, washed with complete medium and PBS, and stained with FCP-1 (5 μM) in DPBS for 45 min. At the last 5-min incubation, a nucleus stain, DRAQ-5 (5 μM), was added to the solution mixture. Without washing, the cells were imaged by confocal fluorescence microscopy.

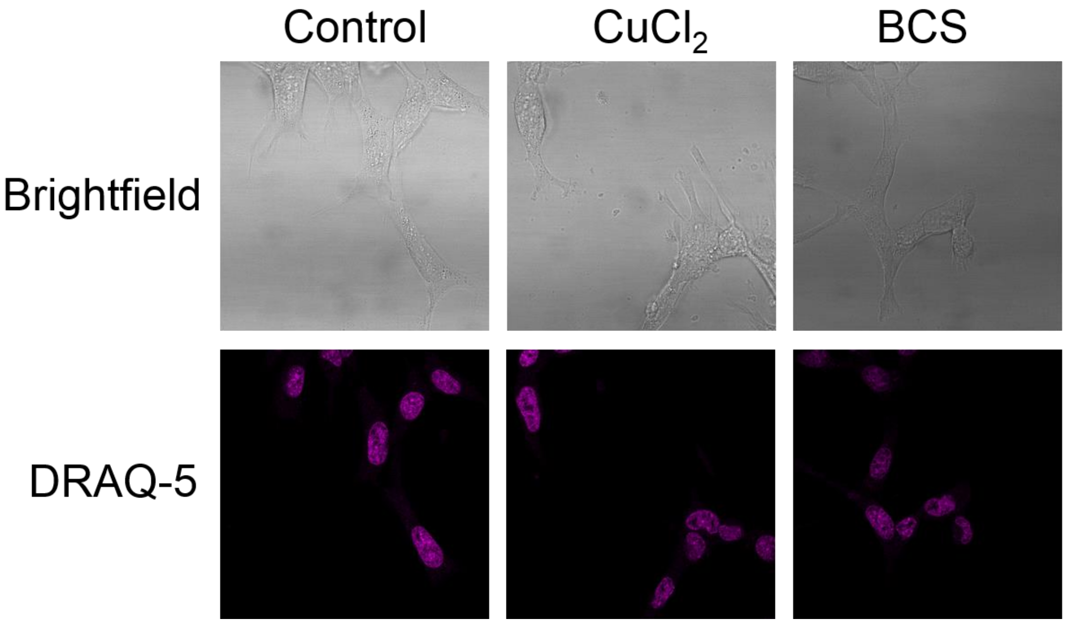


**Fig. S15.** Confocal microscopy images of *Ctr1*$^{-/-}$ MEFs pretreated with solvent vehicle, $CuCl_2$ (100 μM) or BCS (500 μM) in complete medium for 8 h, washed with PBS and stained with FCP-1 (5 μM) in DPBS for 45 min. At the last 5-min incubation, a nucleus stain, DRAQ-5 (5 μM), was added to the solution mixture. Without washing, the cells were imaged by confocal fluorescence microscopy.

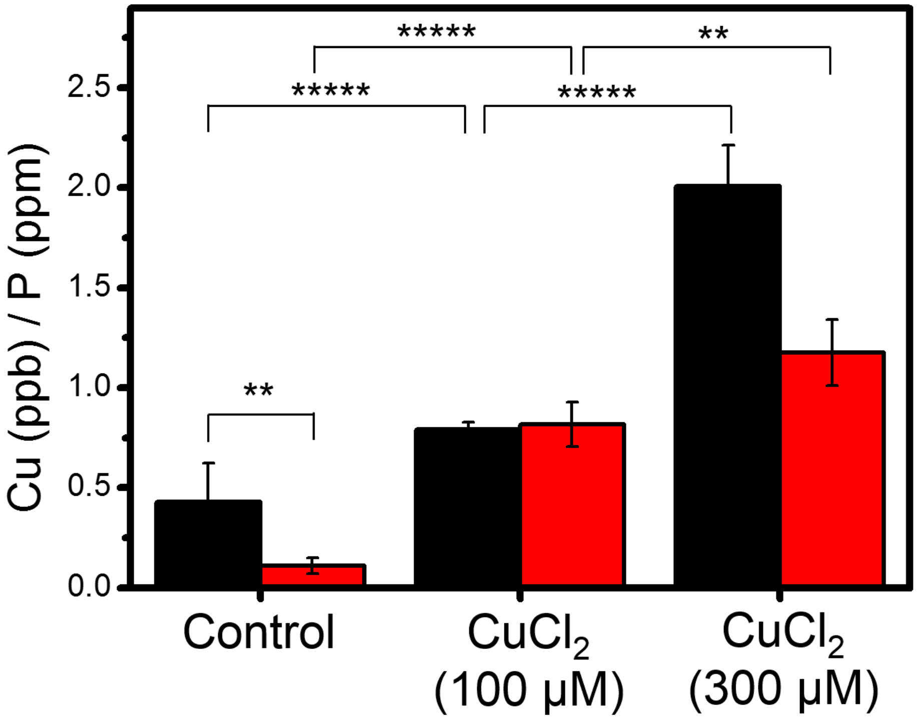


**Fig. S16.** ICP-MS measurement to determine total cellular $^{63}$Cu levels (with normalization of different cell numbers by total cellular $^{31}$P level) in *Ctr1*$^{+/+}$ MEFs (black) and *Ctr1*$^{-/-}$ MEFs (red) treated with solvent control or $CuCl_2$ in complete medium for 8 h. Error bars denote SD ($n$ = 5). **$p$ < 0.01 and *****$p$ < 0.00001. Increases in total Cu levels in both *Ctr1*$^{+/+}$ and *Ctr1*$^{-/-}$ MEFs upon $CuCl_2$ treatment were measured despite the observation that labile Cu(I) pools were less altered in $CuCl_2$-treated *Ctr1*$^{-/-}$ MEFs cells relative to wildtype, suggesting a more complex interplay between total and labile copper homeostasis in the *Ctr1*$^{-/-}$ MEFs models upon copper supplementation.

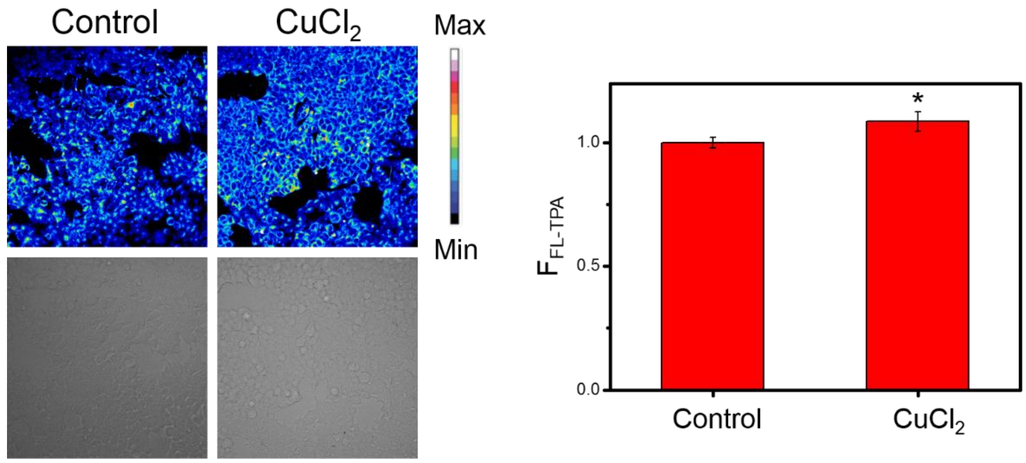


**Fig. S17.** Confocal microscopy images of HEK 293T cells pretreated with solvent vehicle or $CuCl_2$ (25 μM) in complete medium for 18 h, washed with PBS, stained with FL-TPA (5 μM) in DPBS for 45 min and imaged without washing. $\lambda_{ex}$ = 488 nm.

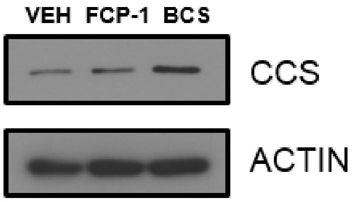


**Fig. S18.** Immunoblot detection of CCS or ACTIN in MEFs treated with vehicle (VEH), 5 µM FCP-1, or 0.5 mM BCS.

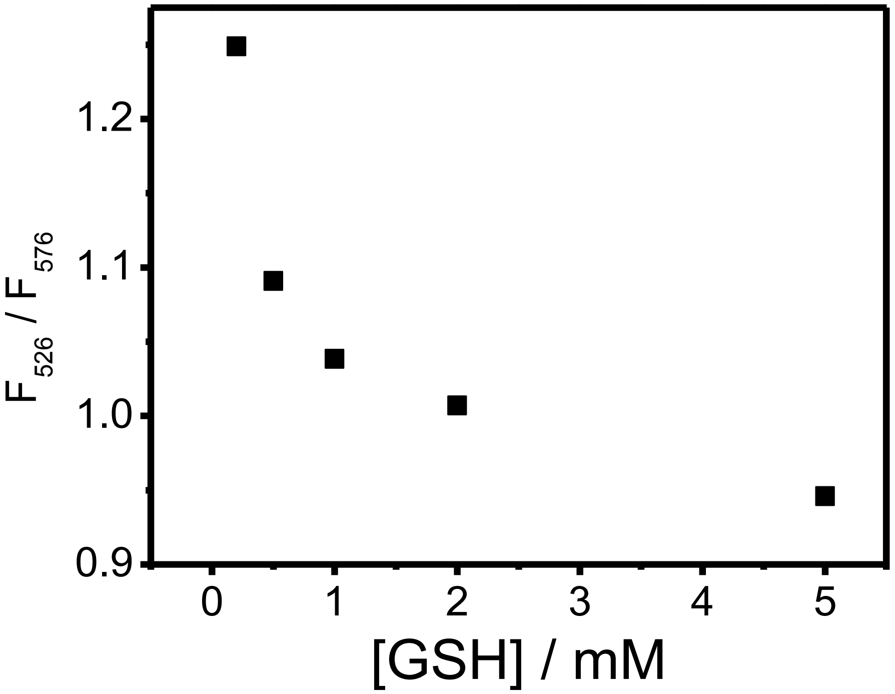


**Fig. S19.** $F_{526}$ / $F_{576}$ of FCP-1 (5 μM) and Cu(I) (10 μM) in aqueous buffer solution (PBS containing 40 vol% PEG-400) with different concentrations of GSH after incubation for 15 min. $\lambda_{ex}$ = 458 nm.

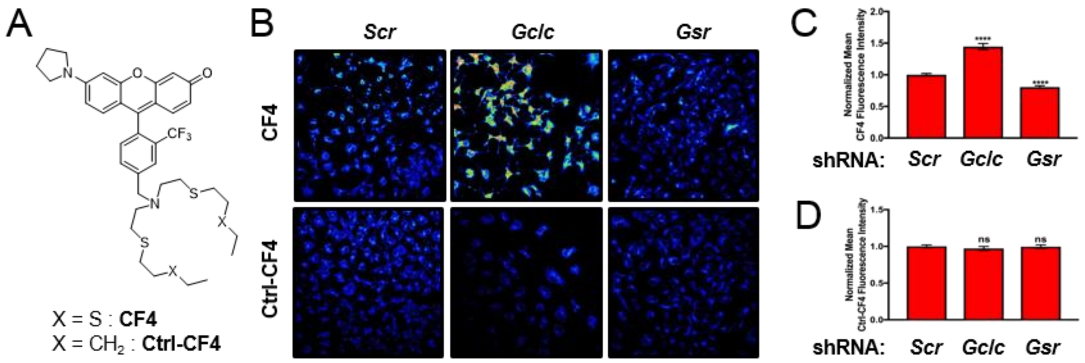


**Fig. S20.** (A) Chemical structures of CF4 and Ctrl-CF4. (B) Representative live cell images of MEFs stably expressing non-targeting control shRNA (*Scr*), *Gclc* shRNA, or *Gsr* shRNA stained by CF4 or Ctrl-CF4. Quantification of mean (C) CF4 or (D) Ctrl-CF4 fluorescence intensity ± s.e.m. from MEFs stably expressing *Scr*, *Gclc* shRNA, or *Gsr* shRNA. Results were compared using a one-way ANOVA followed by a Dunnett's multi-comparisons test. Data are from analysis of 90 or more individual cells (n≥90). ****, $p < 0.0001$.

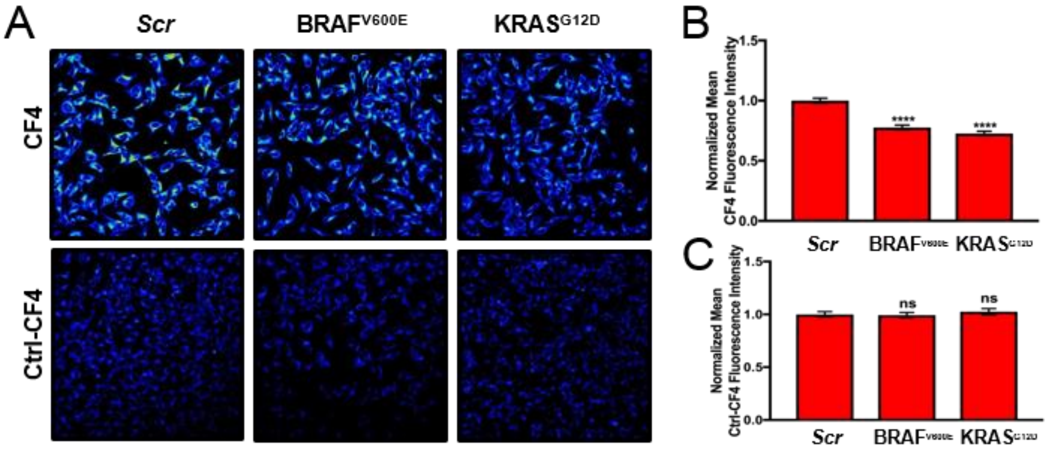


**Fig. S21.** (A) Representative live cell images of MEFs stably expressing *Scr*, *BRAF*$^{V600E}$ cDNA, or *KRAS*$^{G12D}$ cDNA stained by CF4 or Ctrl-CF4. Quantification of mean (B) CF4 or (C) Ctrl-CF4 fluorescence intensity ± s.e.m. from the stained cells. Results were compared using a one-way ANOVA followed by a Dunnett's multi-comparisons test. Data are from analysis of 90 or more individual cells ($n \geq 90$). ****, $p < 0.0001$.

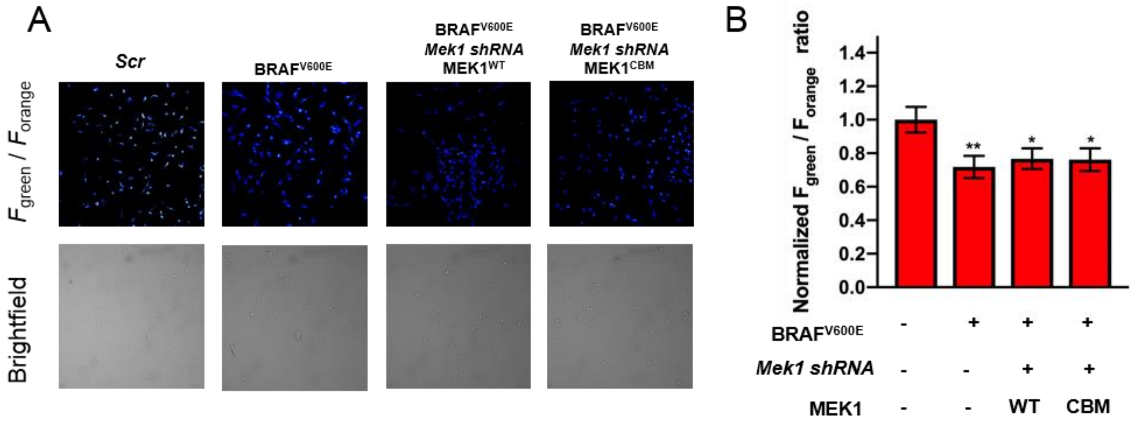


**Fig. S22.** (A) Representative live cell images of MEFs stably expressing *Scr*, *BRAF$^{V600E}$* cDNA, *BRAF$^{V600E}$* cDNA and *Mek1shRNA* with RNA-mediated-interference-resistance wild-type MEK1 *cDNA* or *MEK1$^{CBM}$* cDNA stained by FCP-1. (B) Quantification of mean FCP-1 fluorescence emission ($F_{green}$/$F_{orange}$) ± s.e.m. from stably expressing *Scr*, *BRAF$^{V600E}$* cDNA, *BRAF$^{V600E}$* cDNA and *Mek1shRNA* with RNA-mediated-interference-resistance wild-type MEK1 *cDNA* or *MEK1$^{CBM}$* cDNA Results were compared using a one-way ANOVA followed by a Dunnett's multi-comparisons test. Data are from analysis of 90 individual cells (n=90).
